# Exposure of negative-sense viral RNA in the cytoplasm initiates innate immunity to West Nile virus

**DOI:** 10.1101/2024.06.07.597966

**Authors:** Emmanuelle Genoyer, Jonathan Wilson, Joshua M. Ames, Caleb Stokes, Dante Moreno, Noa Etzyon, Andrew Oberst, Michael Gale

**Affiliations:** Department of Immunology, University of Washington, Seattle WA, USA; Department of Pediatrics, University of Washington/Seattle Children’s Hospital, Seattle WA, USA; Center for Innate Immunity and Immune Disease, University of Washington, Seattle WA, USA

## Abstract

For many RNA viruses, immunity is triggered when RIG-I-like receptors (RLRs) detect viral RNA. However, only a minority of infected cells undergo innate immune activation. By examining these “first responder” cells during West Nile virus infection, we found that specific accumulation of anti- genomic negative-sense viral RNA (-vRNA) underlies innate immune activation and that RIG-I preferentially interacts with -vRNA. However, flaviviruses sequester -vRNA into membrane-bound replication compartments away from cytosolic sensors. We found that single-stranded -vRNA accumulates outside of replication compartments in “first responder” cells, rendering it accessible to RLRs. Exposure of this -vRNA occurs at late timepoints of infection, is linked to viral assembly, and depends on the expression of viral structural proteins. These findings reveal that while most infected cells replicate high levels of vRNA, release of -vRNA from replication compartments during assembly occurs at low frequency and is critical for initiation of innate immunity during flavivirus infection.

## INTRODUCTION

Flaviviruses are a global threat to public health, placing billions at risk of infection and causing over 400 million disease cases per year worldwide^1^. The emerging neurotropic flavivirus West Nile virus (WNV) is the leading cause of arboviral infections in the United States with recurrent annual outbreaks and poses an ever-expanding hazard as its vector range increases^2^. During WNV infection, the initiation of an antiviral innate immune response is critical for viral clearance and engagement of the adaptive immune response^3^. Previous studies form our lab and others have shown that the RIG-I-like receptor (RLR) family of pathogen recognition receptors are critical for initiating the antiviral response to WNV infection and programming the adaptive immune response both in vitro and in vivo^4–7^.

RLRs recognize pathogen associated molecular patterns (PAMPs) in viral RNA in the cytoplasm of infected cells including 5’ tri-phosphate (ppp) RNA, double-stranded RNA, and sequence specific motifs within RNA^8^. Recognition of PAMP RNA initiates a signaling cascade that leads to the phosphorylation of the transcription factor IRF3 with activation of NF-κB and other transcription factors that direct the induction of target genes including interferons, immune-modulatory cytokines, and chemokines. Following production and signaling by types I and III IFN, hundreds of interferon stimulated genes (ISGs) are expressed. Specific ISG products mediate antiviral actions to limit virus replication and spread^9^. Both RIG-I and MDA5 have been implicated in recognition of WNV RNA, with RIG-I playing an early role in detection^4^. However, the type of RNA that is recognized by the RLRs during WNV infection remains unknown. WNV genomic RNA has a 5’ cap-1 structure (m^7^GpppAm)^10^ which prevents RIG-I from biding to RNA by steric hinderance^11,12^. Therefore, it has been hypothesized that uncapped replication intermediates serve as PAMP RNA during infection.

However, flaviviruses, like most positive-strand RNA viruses, sequester their RNA into replication compartments (RCs) that are formed within the endoplasmic reticulum. These RCs serve as sites of viral RNA replication and their enclosure within membranes is thought to facilitate evasion of replication intermediate viral RNA from being sensed by cytoplasmic RNA sensors^13–15^. The hypothesis that WNV evades innate immune activation by passive sequestration of viral PAMPs away from sensors is bolstered by evidence that the virus largely does not encode antagonists of the RLR pathway^16^. Thus, a fundamental question in innate immune programming against WNV and many other positive-sense RNA viruses is how RLRs gain access to viral RNAs to initiate signaling cascades.

An increasing interest in single cell biology has revealed that viral replication and viral RNA accumulation within infected cells during flavivirus infection is extremely heterogeneous^17–20^. Further, it is increasingly apparent that innate immune activation is initiated by a small subset of infected cells, termed “first responder cells”. This phenomenon has been observed during infection with WNV and Zika virus, a related flavivirus^18,20^, as well as during infection by many other viruses including influenza A virus^21,22^, herpes simplex virus ^23^, and encephalomyocarditis virus^24^. By defining the features of viral replication required for innate immune activation in first responder cells we can better understand viral vulnerabilities to host cell detection.

Here we identified and examined these first responder cells during WNV infection and found that they harbor increased levels of single-stranded negative-sense viral RNA (-vRNA). This -vRNA is sensed by and preferentially interacts with RIG-I during infection to drive innate immune activation. Intriguingly, we found that single-stranded -vRNA becomes exposed in the cytoplasm of first responder cells and that this exposure occurs concurrent with viral assembly and the expression of viral structural proteins.

## RESULTS

### Specific accumulation of -vRNA in innate immune activated cells

During viral infections, only a small minority of cells initiate an antiviral innate immune response^18,23–25^. Indeed, during West Nile virus infection in cultured cells there is a logarithmic amplification of viral RNA (**Fig 1A**) followed by induction of interferon beta (IFNΒ) beginning at 18 hours post infection (hpi) (**Fig 1B**). However, when all cells have high levels of viral RNA, only a subset have translocated IRF3 to the nucleus at early timepoints of infection (**Fig 1C-D**), a hallmark of RLR-driven innate immune activation. We found that this heterogeneous response is maintained during infections of different cell types (**Fig 1E-F**) and while percentage of early responding cells changes, these levels are similar to levels reported by other groups for WNV^18^ and Zika virus^20^, another member of the flavivirus family.

**Figure 1.**
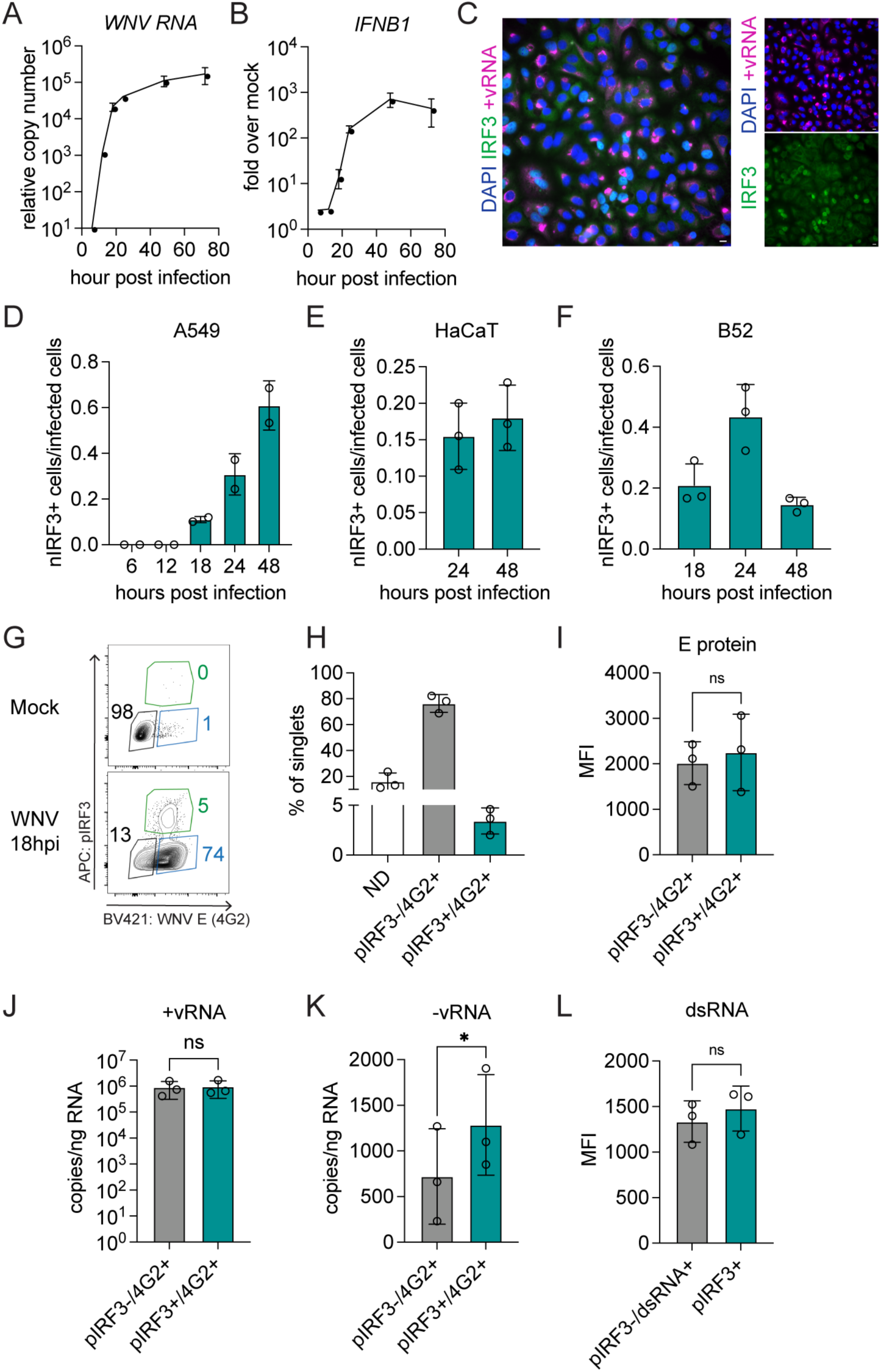
Specific accumulation of -vRNA in first responder cells. A549 cells infected with WNV-TX MOI 1.5 were analyzed by RT-qPCR for **(A)** WNV RNA and **(B)** *IFNB1* mRNA at indicated timepoints. Data is graphed as mean ± SEM of 2-4 experiments. **(C)** Infected cells were subjected to RNA-FISH for +vRNA (magenta) combined with immunofluorescence (green), nuclei counterstained with DAPI (blue). Widefield 60X image, scale bar = 10um. Representative of three independent experiments. Ratio of infected cells with nuclear IRF3 in **(D)** bronchial epithelial A549 cells, **(E)** keratinocyte HaCaT cells and (**F)** skin fibroblast B52 cells. **(G)** Representative flow plot and gating strategy of A549 cells infected with WNV-TX MOI 1.5 18h analyzed for innate immune activation by phospho-IRF3 and infection by presence of WNV E (4G2), pIRF3+/4G2+ population boxed in green, pIRF3-/4G2+ population boxed in blue, non-detectable infection (ND) boxed in black. **(H)** Relative frequency of populations. **(I)** Viral protein abundance (WNV E) in subpopulations by geometric mean fluorescence intensity (MFI). Cells were sorted and strand-specific RT-qPCR was used to quantify **(J)** +vRNA and **(K)** -vRNA in sorted populations. **(L)** dsRNA levels were measured by geometric mean fluorescence intensity in each subpopulation. Data is graphed as mean ± SD with individual experiments represented. * = p<0.05, ns = not significant by paired t-test.

To determine how accumulation of viral RNA corresponded to innate immune activation in first responder cells, we performed cell sorting based on immunofluorescence intensity for WNV antigen and IRF3 phosphorylation at 18 hours post infection, the earliest timepoint at which we detect innate immune activation in infected cells (**Fig 1G-H**). We found no difference in the levels of viral envelope protein expression by mean fluorescence intensity when comparing infected non-responsive (4G2+/pIRF3-) to infected responsive (4G2+/pIRF3+) cells. (**Fig 1I**). We then extracted RNA to measure strand-specific viral RNA levels. While responsive cells were not enriched for positive-sense viral RNA (+vRNA) (**Fig 1J**), they harbored increased levels of negative-sense replication intermediate viral RNA (-vRNA) (**Fig 1K**). Because double-stranded RNA (dsRNA) is thought to be a viral PAMP during infection^26^, we evaluated levels of dsRNA within individual cell types by mean fluorescence intensity using the anti-dsRNA antibody J2. Our data demonstrate no specific enrichment of dsRNA in the first responder cells (**Fig 1L**). Therefore, first responder cells specifically carry higher levels of -vRNA, suggesting that RLR engagement specifically by negative-sense viral RNA is the key step leading to activation of innate immunity in first responder cells.

### -vRNA is a preferential substrate for RIG-I during infection

To determine if -vRNA is a preferential substrate for RLRs during infection, we evaluated WNV-indued innate immune activation in A549 epithelial cells. While WNV can be recognized *in vivo* by RIG-I, MDA5, and TLR3, we chose to use the A549 infection model in which only RIG-I participates in recognition of WNV (**Figure S1**)^27^. To determine the species of viral RNA that are recognized and bound by RIG-I during WNV infection, we performed reversible formaldehyde crosslinking followed by immunoprecipitation of RIG-I in A549-IRF3-knockout (KO) cells. IRF3KO cells allow for immunoprecipitation solely of endogenous basal levels of RIG-I involved in original detection of viral PAMP, without RIG-I expression induction by IFN. Our approach allowed for cross-linking prior to immunoprecipitation to prevent reassortment of RNA-protein interactions after cell lysis, while still allowing for PCR following crosslink reversal^28^. We next conducted strand-specific qPCR analyses of immunoprecipitated products. We found -vRNA to be clearly enriched compared to +vRNA in RIG-I immunoprecipitation over serum control immunoprecipitation products (**Fig 2A-B**). This outcome indicates that -vRNA is a preferential substrate for RIG-I during WNV infection.

**Figure 2.**
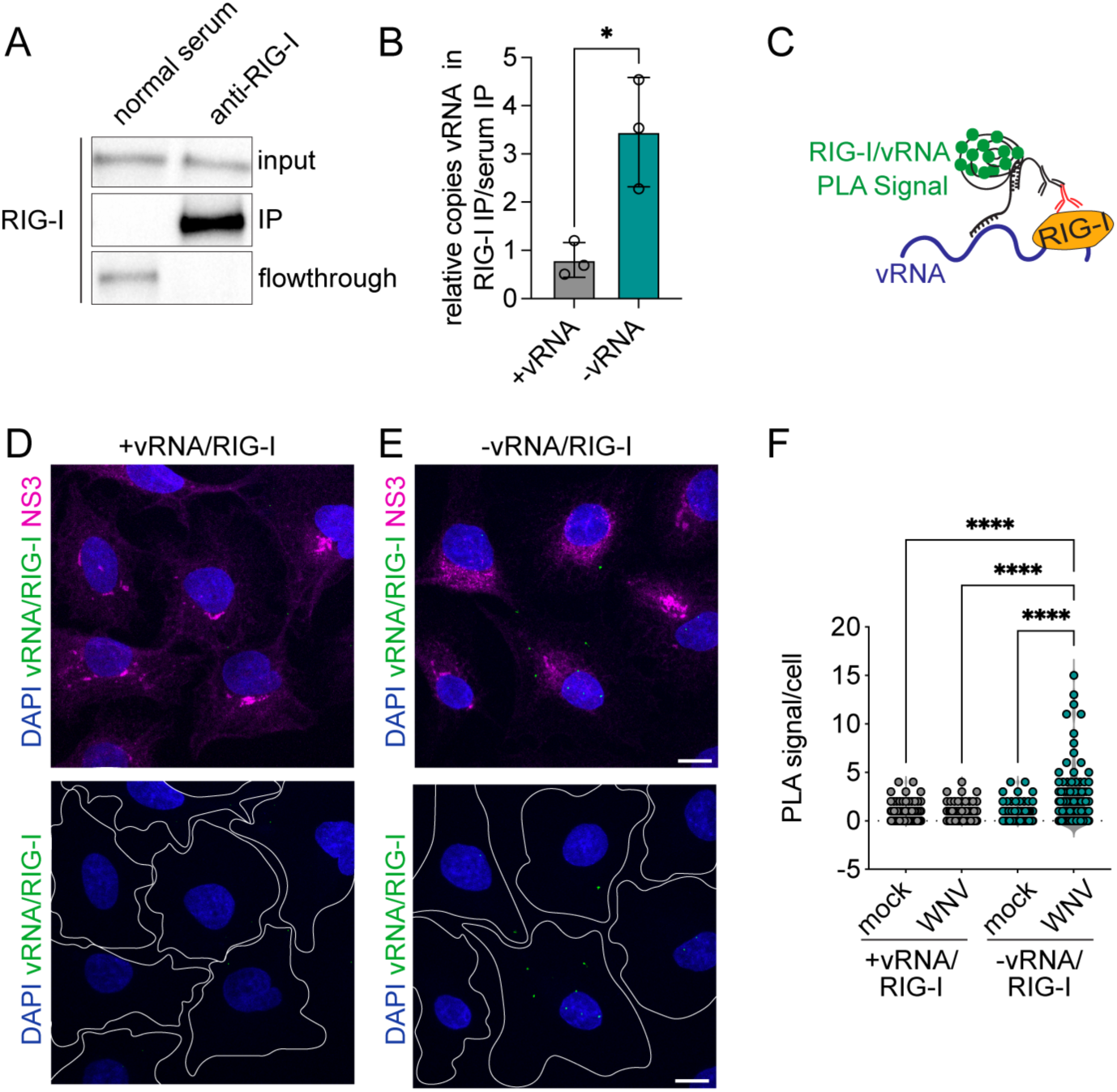
RIG-I preferentially interacts with WNV negative-sense vRNA. **(A)** A549 IRF3KO cells infected with WNV-TX for 24h followed by formaldehyde crosslinked immunoprecipitation of RIG-I or control (normal rabbit serum). Western blot of RIG-I to assess immunoprecipitation. **(B)** RNA was extracted from immunoprecipitations and assayed for vRNA by strand-specific RT-qPCR. **(C)** Schematic of RIG-I/vRNA proximity ligation assay with probes specific to -vRNA and +vRNA. **(D-E)** A549-RML cells were infected with WNV-TX MOI 1.5 for 18h and subjected to vRNA/RIG-I PLA (green) and immunofluorescence was used to detect viral infection by NS3 (magenta), nuclei were counterstained with DAPI (blue). Cells outlined in white in bottom panel. Images are confocal 60x z-stack max-projections, representative of three independent experiments. Scale bar = 10um. **(E)** PLA puncta per cell was quantified for mock and WNV infected cells. Individual cell values are graphed overlaid onto violin plot, data is a sum of three independent experiments. **** = p<0.0001 by Kruskal-Wallis test with Dunn’s multiple comparisons test.

To further characterize interactions of RIG-I and vRNA *in situ* we performed a modified proximity ligation assay (PLA) to detect RNA-protein interactions (RNA-PLA) (**Fig 2C**)^29^. Because basal levels of RIG-I are difficult to visualize, we took advantage of A549-RML cells that express slightly elevated levels of RLRs, including RIG-I, but retain the same kinetics of innate immune activation and viral replication^30^ (**Fig S2**). RNA-PLA to detect interaction between +vRNA and RIG-I at 18 hpi showed no difference in signal between mock and infected cells. In contrast, when -vRNA/RIG-I contacts were assayed, PLA signal significantly increased above the level in observed in mock infected cells (**Fig 2D-F**). Together, these results indicate that RIG-I preferentially interacts with -vRNA within infected cells.

### Distinct spatiotemporal accumulation of +vRNA and -vRNA during infection

RT-qPCR quantification indicates that levels of +vRNA and -vRNA increase logarithmically during acute WNV replication, with levels of +vRNA existing at 100-1000-fold higher levels than - vRNA throughout infection (**Fig 3A-B**)^31^. During flavivirus replication, the replication-intermediate vRNA (-vRNA) is thought to reside in replication compartments where it serves as a template for +vRNA transcription^14,32^. To further evaluate vRNA interactions with RLRs during infection, we characterized the accumulation and distribution of positive- and negative-sense vRNA over time using RNAscope. RNAscope employs strand-specific oligos followed by a series of amplifications to visualize individual RNAs within cells^33^. Due to the nature of probe binding, only single stranded RNA is normally detected. However, we found that denaturation of dsRNA prior to probe binding revealed all RNA within a cell, including that which was previously double-stranded (**Fig 3C**). The denaturation of dsRNA can then be validated by a decrease in the detection of dsRNA by the dsRNA-specific antibody J2 (**Fig 3D**).

**Figure 3.**
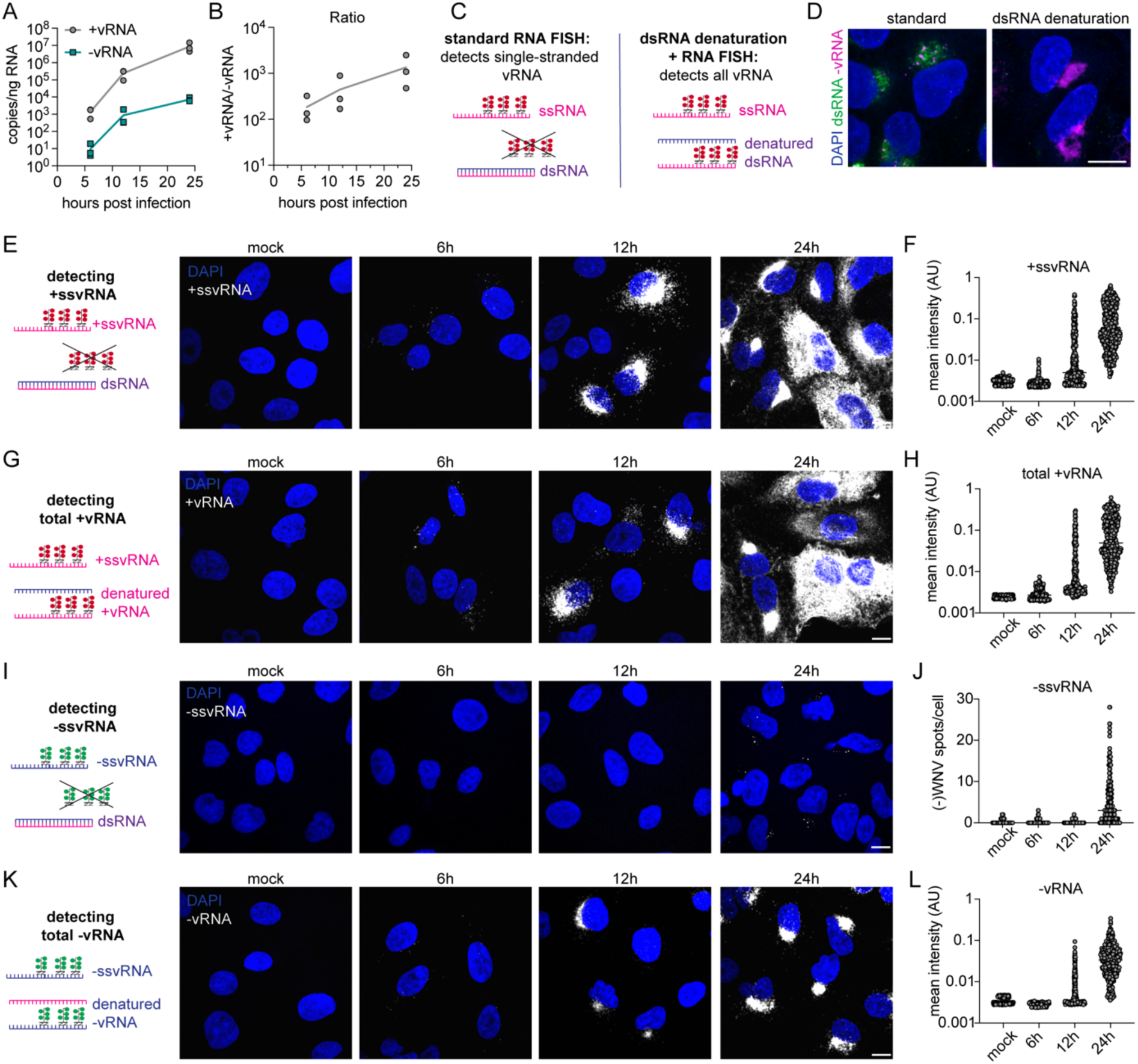
Distinct spatiotemporal accumulation of positive and negative sense vRNA during infection. **(A)** Strand-specific RT-qPCR to detect +/-vRNA at indicated timepoints in A549 cells infected with WNV-TX. **(B)** Ratio of +/-vRNA at indicated timepoints. Data is graphed as mean of three individual experiments with individual experiments represented. **(C)** Experimental approach to detect single stranded vRNA and total vRNA by RNAscope. **(D)** A549 cells infected for 24h with WNV-TX MOI 1.5 were subjected to immunofluorescence to detect dsRNA (green) followed by RNAscope to detect -vRNA (magenta) without (standard) or with (dsRNA denaturation) RNA denaturation prior to probe binding. Nuclei are counterstained with DAPI (blue). **(E)** RNAscope on WNV infected A549 cells at indicated timepoints to detect +ssvRNA (white) **(F)** Quantification of intensity of +ssvRNA in individual cells. **(G)** RNAscope on WNV infected A549 cells at indicated timepoints to detect total +vRNA with dsRNA denaturation **(H)** Quantification of intensity of total +vRNA in individual cells. **(I)** RNAscope on WNV infected A549 cells at indicated timepoints to detect -ssvRNA **(J)** Quantification of - ssvRNA puncta in individual cells. **(K)** RNAscope on WNV infected A549 cells at indicated timepoints to detect total -vRNA with dsRNA denaturation **(L)** Quantification of intensity of total -vRNA in individual cells. Images are confocal 60x z-stack max-projections, representative of three independent experiments, scale bar = 10um. Individual cell values are graphed, data is a sum of three independent experiments with line at mean. AU = arbitrary units.

Employing both methods, we visualized and quantified +vRNA within infected cells across an acute infection time course (**Fig 3E-H**). At early timepoints (6 hpi), single-stranded +vRNA (+ssvRNA) is punctate and sparse within infected cells, eventually coalescing in the perinuclear region (12 hpi) and expanding throughout the cytoplasm at late timepoints (24 hpi). Single-stranded +vRNA (+ssvRNA) is detected at all timepoints (**Fig 3E-F**). Assessment of total +vRNA upon denaturation showed similar distribution patterns across time as -ssvRNA (**Fig 3G-H**). Quantification of intensity of +ssvRNA and total +vRNA within individual cells highlights the heterogeneity of vRNA accumulation within cells (**Fig 3F, 3H**). Quantification of +vRNA levels and localization patterns were similar between ssvRNA and total vRNA, indicating that the vast majority of +vRNA exists in a single-stranded form during WNV infection.

In contrast, single-stranded -vRNA (-ssvRNA) was undetectable at 6 and 12 hpi. Importantly, discrete puncta of -ssvRNA were detectable at 24 hpi (**Fig 3I-J**). When applying RNAscope after dsRNA denaturation, total -vRNA was detectable at all timepoints (**Fig 3K-L**). At 6 hpi, -vRNA localized to puncta similar to distribution of +vRNA, likely marking early replication compartments. By 12 hpi -vRNA coalesced perinuclearly similar to +vRNA, and remained similar in distribution at 24 hpi. While intensity overall was lower than +vRNA, quantification of total - vRNA within single cells maintained the general trend of +vRNA accumulation. These measurements indicate that a vast majority of -vRNA exists as double-stranded throughout the course of infection. Critically, it is the appearance of -ssvRNA at late time points during infection that coincide with innate immune activation (**Fig 1B**).

### -ssvRNA is localized outside of replication compartments

To visualize the location of -ssvRNA in relation to viral replication complexes (RCs) we performed RNAscope coupled with immunofluorescence for the viral protein NS3. NS3 is the bifunctional viral protease/helicase that participates in viral replication and resides within viral RCs^15,34^. Interestingly, confocal imaging revealed that the -ssvRNA was localized discretely from NS3, indicating that -ssvRNA was present outside of replication compartments (**Fig 4A-B**). Because the flavivirus RC functions to sequester viral RNA away from detection by cytosolic PRRs, the cytosolic -ssvRNA might engage innate immune signaling due to its mislocalization outside of the RC.

**Figure 4.**
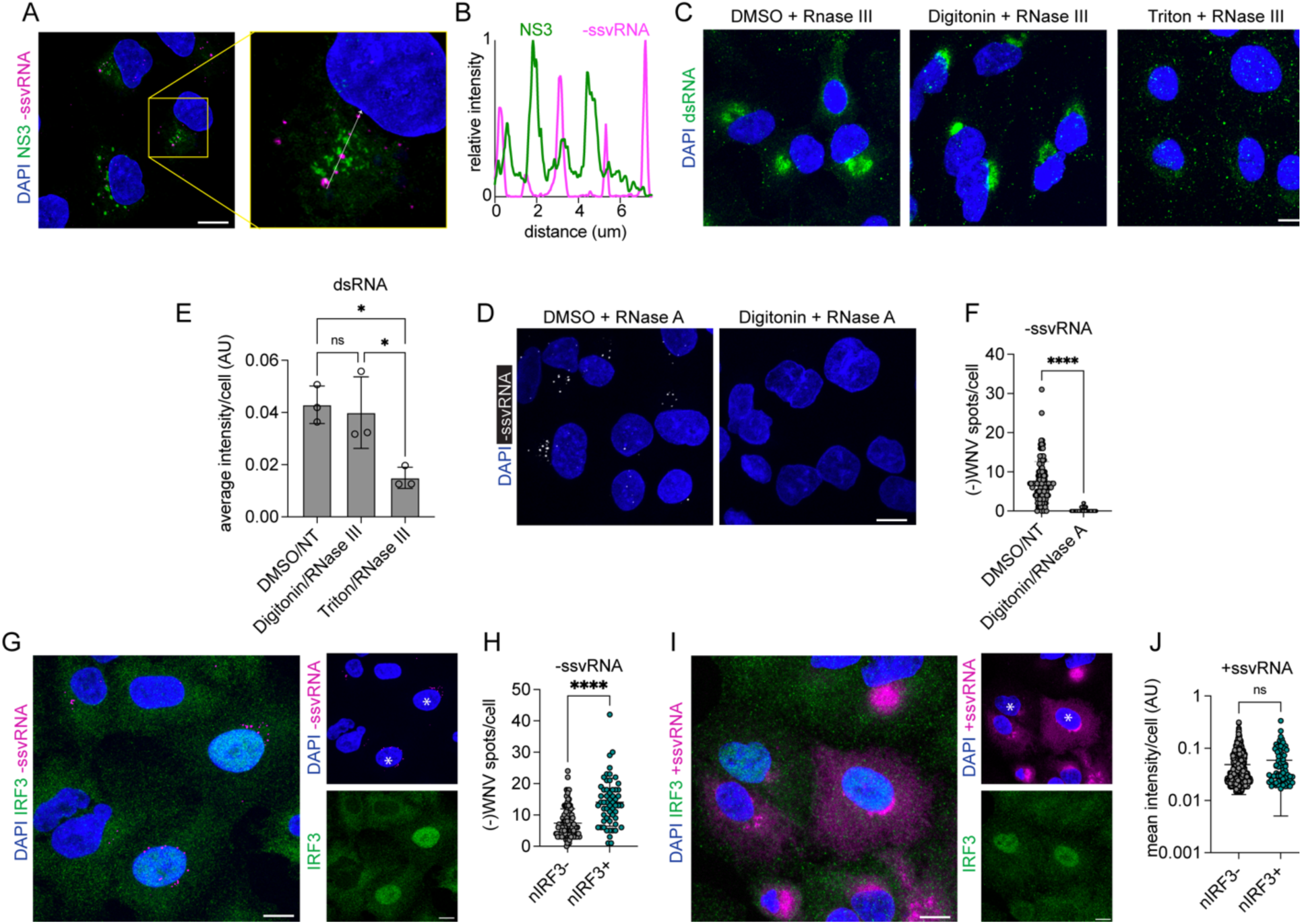
-ssvRNA is localized outside of replication compartments and links with innate immune induction. **(A)** A549 cells were infected with WNV-TX MOI 1.5 for 24h followed by combined immunofluorescence for NS3 (green) and RNAscope to detect -ssvRNA (magenta), nuclei counterstained with DAPI (blue); confocal 60x image z-stack max projection with lightening deconvolution. Image is representative of three independent experiments. **(B)** Plot profile of arrow in inset in (A) of normalized relative intensities of NS3 (green) and -ssvRNA (magenta). A549 cells were infected with WNV-TX MOI 1.5 for 24h followed by permeabilization and RNase treatment as indicated prior to fixation and subjected to **(C)** immunofluorescence to detect dsRNA (green), nuclei counterstained with DAPI (blue). **(D)** Mean intensity of dsRNA was calculated per cell, data represent average cell intensity from three independent experiments, graphed as mean ± SD, **** = p<0.0001 by one-way ANOVA with Tukey’s multiple comparison. **(E)** RNAscope to detect -ssvRNA (white), nuclei counterstained with DAPI (blue). **(F)** -ssvRNA puncta were quantified per cell, data represent individual cells from three independent experiments with line at mean, **** = p<0.0001 by Mann-Whitney U-test. **(G)** A549 cells were infected with WNV-TX MOI 1.5 for 18h then subjected to combined immunofluorescence for IRF3 (green) and RNAscope to detect -ssvRNA (magenta), nuclei counterstained with DAPI (blue). White asterisk indicates cell with nuclear IRF3. **(H)** -ssvRNA puncta were quantified per cell. **(I)** A549 cells were infected with WNV-TX MOI 1.5 for 18h then subjected to combined immunofluorescence for IRF3 (green) and RNA-FISH to detect +ssvRNA (magenta), nuclei counterstained with DAPI (blue). White asterisk indicates cell with nuclear IRF3. **(J)** mean intensity of +ssvRNA was calculated per cell. Data represent individual cells from three independent experiments, with line at mean **** = p<0.0001, ns= not significant by Mann-Whitney U-test. 60x confocal z-stack max projection images are representative of three independent experiments, scale bar = 10um.

To further interrogate the location of the single-stranded -vRNA in relation to RCs, we used a biochemical approach to assess -vRNA vulnerability to RNase digestion upon selective membrane permeabilization. Treatment of cells with a mild detergent like digitonin results in the permeabilization of the plasma membrane but not intracellular membranes. Viral RNA residing in replication compartments within the ER membrane thus is not accessible with digitonin permeabilization. The use of stronger detergents such as Triton X-100 is required to permeabilize ER membranes^35^. We first confirmed this specificity by treating infected cells with either digitonin or Triton X-100, followed by exposure to RNase III, which digests dsRNA. In cells treated with digitonin, there was no significant change in dsRNA signal, indicating viral RNA remained protected within intact replication compartments. In contrast, the dsRNA signal was significantly decreased upon treatment with Triton X-100 and RNaseIII (**Fig 4C-D**). However, digitonin alone followed by RNase A treatment, which digests ssRNA, was sufficient to completely abolish the - ssvRNA signal (**Fig 4E-F**). This outcome indicates that these -ssvRNAs are indeed located outside of replication compartments and biochemically confirms that they are single-stranded.

### -ssvRNA accumulation links with innate immune activation

During WNV infection we observed that the accumulation of -ssvRNA is heterogeneous amongst cells with some cells having few puncta to some cells having upwards of 30 puncta by 24 hours post infection (**Fig 3J**). To determine if the differences in accumulation of accessible - ssvRNA links with differences in innate immune activation status, we performed RNAscope coupled with immunofluorescence to define the subcellular localization and activation state of IRF3. We found that -ssvRNA puncta are significantly increased in nuclear IRF3 (nIRF3) -positive cells (**Fig 4G-H**). In contrast, there is no significant difference in mean fluorescence intensity of +ssvRNA between nIRF3-positive and nIRF3-negative cells (**Fig 4I-J**). This supports the conclusion that -ssvRNA is the preferential PAMP for RIG-I.

### RIG-I levels determine threshold of -ssvRNA required for innate immune activation

Heterogeneity in basal expression of pattern recognition receptors is thought to contribute to differential levels and rates of immune activation^18,36^. RIG-I itself is an interferon-stimulated gene (ISG), meaning that production of interferon by first responder cells during infection may position bystander cells to respond to infection more rapidly owing to increased levels of RIG-I. To test whether levels of RIG-I may control the threshold of -ssvRNA required to activate an innate immune response, we generated A549 cells that stably over-expressed RIG-I (termed RIG-I OE) (**Fig S3A**). Overexpression of RIG-I did not affect early kinetics of viral infection, nor did it shift the kinetics of innate immune activation (**Fig S3B-C**), however levels of *IFNB1* were significantly higher once innate immune activation was initiated (**Fig S3C**). When individual cells were examined by flow cytometry, a significantly higher percentage of RIG-I OE cells had phosphorylated IRF3 (**Fig S3D**). When levels of -ssvRNA were examined in individual cells, we found that cells with nuclear IRF3 had significantly higher levels of -ssvRNA than non-responsive cells, recapitulating results found in WT cells (**Fig S3E-F**). When compared to WT cells, overall distribution and levels of -ssvRNA were not different between cell types (**Fig S3G**), but nIRF3-negative (non-responsive) cells had significantly lower levels of -ssvRNA (**Fig S3H**). Thus, while increased levels of RIG-I in the cytoplasm do allow cells to initiate innate immune activation with lower levels of accessible, cytosolic PAMP, cells are unable to detect and respond to viral RNA until at least 18 hpi when -ssvRNA becomes exposed within the cytoplasm of infected cells.

### -ssvRNA becomes exposed during viral assembly and is induced by expression of capsid

Because -ssvRNA accumulation occurs at late time points in the course of infection it could be a product of viral particle production. To separate the replication of viral RNA from virion production, we utilized a WNV replicon lacking the structural protein-coding region of the viral genome, which produces vRNA but does not make viral particles **(Fig 5A)**^37^. Attempts to rescue A549 WT cells with replicon were unsuccessful due to the immunostimulatory nature of transfected viral RNA which drives an IRF3-mediaited antiviral response and subsequent cell death. Thus, we used A549-IRF3KO cells bearing WNV replicon to compare levels of -ssvRNA between WNV-replicon and replication-competent WNV for 24 h (**Fig 5B**). Our data demonstrate a striking decrease in the number of -ssvRNA puncta in replicon-bearing cells compared to WNV-infected cells (**Fig 5C**). To confirm the presence of -vRNA in replicon-bearing cells we visualized total -vRNA using dsRNA denaturation prior to probe binding and found increased intensity of - vRNA in replicon-bearing cells (**Fig 5D-E**). Overall, while the A549-IRF3KO cells harboring the WNV replicon had measurable levels of +vRNA and -vRNA, these levels were slightly lower than in WNV-infected A549 IRF3KO cells (**Fig 5F-G**). The difference in -ssvRNA accumulation between replicon and virus-infected cells was recapitulated in Huh7 hepatoma cells either infected with WNV for 24h or harboring the WNV replicon (**Fig 5H-I**).

**Figure 5.**
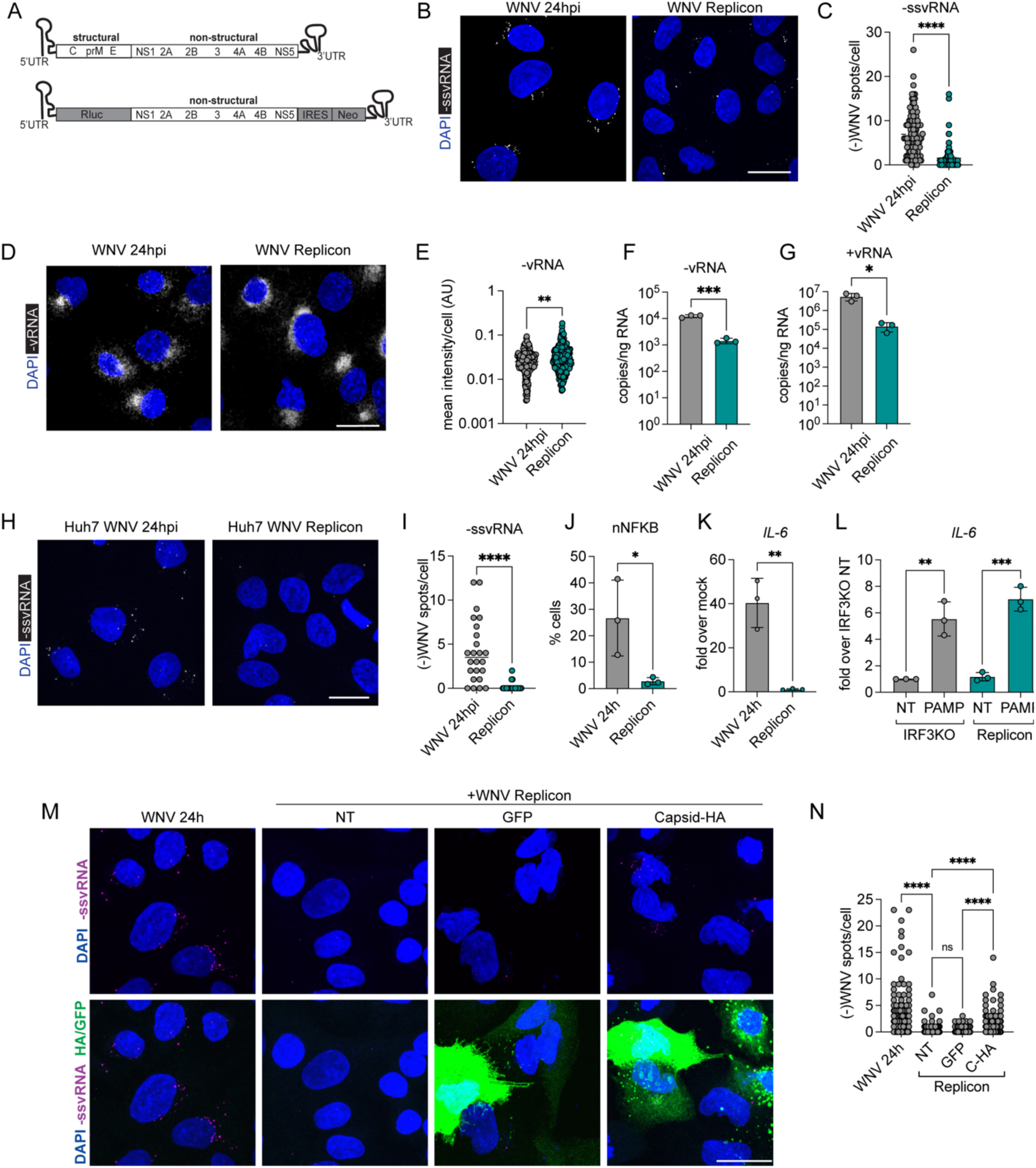
-ssvRNA is exposed during assembly and its absence reduces innate immune activation. **(A)** Schematic of gene arrangement in infectious WNV and WNV replicon. **(B)** A549-IRF3KO cells were infected with WNV-TX for 24h or transfected then selected to harbor a WNV replicon and -ssvRNA was visualized by RNAscope (white), nuclei counterstained with DAPI (blue) **(C)** -ssvRNA puncta were counted per cell, data represent individual cells from three independent experiments, **** = p<0.0001 by Man-Whitney U-test. **(D)** Total -vRNA (white) was visualized by dsRNA denaturation prior to RNAscope probe binding. **(E)** Mean intensity of -vRNA was calculated per cell, data represent individual cells from two independent experiments. Strand-specific RT-qPCR was used to measure **(F)** -vRNA and **(G)** +vRNA levels in WNV infected and WNV replicon-bearing cells, data from three independent experiments graphed as mean ± SD * = p< 0.05, ***=p<0.001 by unpaired t-test **(H)** Huh7 cells were infected with WNV-TX for 24h or transfected then selected to harbor a WNV replicon and -ssvRNA was visualized by RNAscope (white), nuclei counterstained with DAPI (blue) **(I)** -ssvRNA puncta were counted per cell, data represent individual cells from one representative experiment, **** = p<0.0001 by Man-Whitney U-test. **(J)** A549-IRF3KO cells infected with WNV-TX for 24h and A549-IRF3KO replicon-harboring cells were subjected to immunofluorescence to detect total NF-κB. The percentage of cells with nuclear NF-κB were quantified, data is represented as mean ± SD, with individual experiments represented, * = p< 0.05 by unpaired t-test. **(K)** Levels of *IL-6* mRNA relative to mock were measured by RT-qPCR in A549-IRF3KO cells infected with WNV-TX for 24h or harboring replicon, data is graphed as mean ± SD with individual data points plotted. ** = p < 0.01 by unpaired t-test. **(L)** RT-qPCR to detect *IL-6* mRNA levels after transfection of A549-IRF3KO or A549-IRF3KO harboring WNV replicon cells with HCV poly U/UC PAMP RNA for 6 hours, data is graphed as mean ± SD with individual data points plotted, ** = p < 0.01, *** = p<0.001 by unpaired t-test. **(M)** A549-IRF3KO replicon-harboring cells transfected with plasmid expressing GFP or Capsid were subjected to RNAscope for -ssvRNA (magenta) and immunofluorescence to detect C or GFP (green), nuclei counterstained with DAPI (blue); A549-IRF3KO cells were infected with WNV for 24h and A549-IRF3KO replicon-harboring cells were not transfected (NT) as control. **(N)** -ssvRNA puncta were counted per cell, data represent individual cells from three independent experiments, **** = p<0.0001 by Kruskal-Wallis test with Dunn’s multiple comparison test. All images are 60X confocal microscopy, z-stack max projections, scale bar = 20um.

Because we observed an increase in innate immune activation in the presence of cytosolic -ssvRNA, we reasoned that innate immune activation would be decreased in replicon-bearing cells compared to virus-infected cells. RLR activation leads to the downstream phosphorylation, activation, and nuclear translocation of IRF3 and NF-κB^9^. Because our replicon-bearing cells lack IRF3, we used NF-κB activation and target gene transcription as biomarkers of RLR activation. We found that replicon-bearing cells had significantly lower levels of nuclear NF-κB and expression of IL-6 compared to cells infected with WNV for 24 h (**Fig 5J-K**). To confirm that replicon cells could respond to RLR stimulation to direct NF-κB nuclear translocation and IL-6 gene expression, we transfected cells with a potent RIG-I PAMP, an in vitro transcribed region of the polyU/UC region of HCV^38–40^. Upon stimulation of RIG-I, replicon-bearing and infected cells responded with similar activation of NF-κB and induction of IL-6 gene expression (**Fig 5L**). Together, these findings support the notion that cytosolic -ssvRNA accumulates as a by-product of viral particle assembly and confirm that -ssvRNA is the critical PAMP engaging innate immune PRRs.

The reduced -ssvRNA puncta outside replication compartments in replicon-bearing cells might be due to a lack of viral structural proteins in the replicon system. Since the viral capsid protein (C) is known to interact with +vRNA and regulate its packaging, we examined possible C involvement in the accumulation of cytosolic -ssvRNA. WNV replicon-bearing cells were transfected with plasmid expressing HA-tagged capsid protein (C) or GFP as a negative control for 24 hours. We then performed RNAscope for -ssvRNA combined with immunofluorescence to identify transfected cells (**Fig 5M**). We found that complementation with C was sufficient to increase levels of -ssvRNA within cells when compared to non-transfected cells or cells transfected with GFP (**Fig 5N**). This outcome points to a role for capsid protein in either the release of -ssvRNA from replication compartments or for the stabilization of -ssvRNA upon its release from RCs. However, the accumulation of -ssvRNA in C-expressing replicon-bearing cells was consistently lower than levels occurring in WNV-infected cells. This difference may be due to the cellular response to transfection or may reflect a necessity for appropriate stochiometric expression of viral proteins in the release or accumulation of -ssvRNA.

### -ssvRNA accumulates in vivo and correlates with immune activation in neural cell cultures

During natural infections West Nile virus primarily replicates in peripheral myeloid cells. If not controlled in the periphery, WNV can cross the blood brain barrier to establish brain infection that promotes encephalitis^41^. In the brain, replicating WNV is found primarily in neurons, and to a lesser extent in astrocytes and brain microvascular endothelial cells^42^. To test whether -ssvRNA accumulation occurs *in vivo* during infection, we performed RNAscope and immunofluorescence for WNV RNA and protein on brain slices from mice following intracranial infection with WNV. Mice infected intracranially with WNV had extensive viral replication by five days post infection as measured by WNV E protein accumulation in the cortex and hippocampus. We also detected accumulation of -ssvRNA within a subset of infected cells within the cortex (**Fig 6A-B**). Quantification of individual puncta within infected cells revealed extreme heterogeneity in accumulation of -ssvRNA, with the majority of infected cells accumulating little to no -ssvRNA, while a limited number of cells contained more than 60 puncta, consistent with first responder cells (**Fig 6C**). Thus, -ssvRNA is present in vivo.

**Figure 6.**
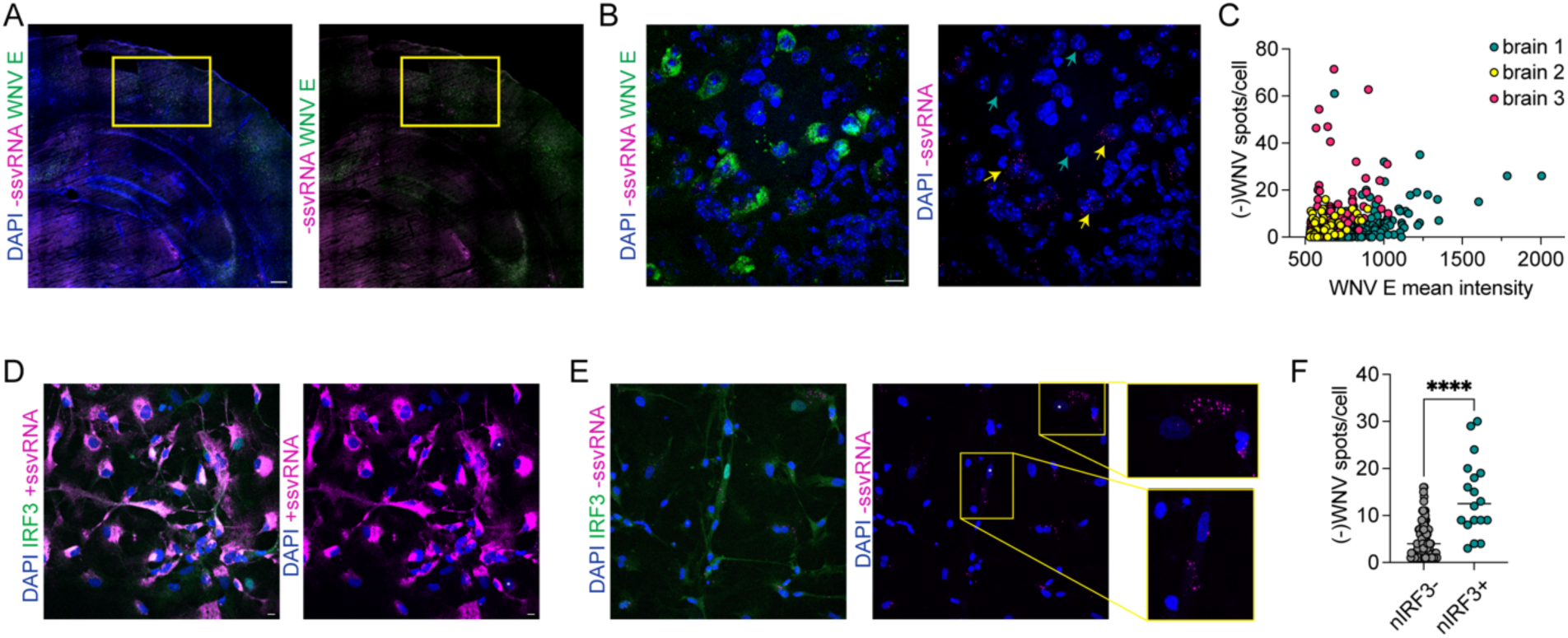
-ssvRNA accumulates in infected brains and correlates with innate immune activation in primary neuronal cultures. **(A)** -ssvRNA by RNAscope (magenta) with immunofluorescence to detect WNV E (green) in mouse brain tissue sections, nuclei counterstained with DAPI (blue), 5 days post intracranial infection with WNV. 20X magnification tile-stitched, scale bar = 200um. Cortex boxed in yellow. **(B)** Confocal 60X, z-stack maximum projections highlighting infected cells in the cortex with high levels of -ssvRNA (yellow arrows) and low/no -ssvRNA (teal arrows), scale bar = 10um. **(C)** -ssvRNA puncta were counted per infected cell and plotted against mean intensity values of WNV E. **(D)** Differentiated induced neural progenitor cells were infected with WNV at MOI 2 for 24h then subjected to combined immunofluorescence and RNAscope to detect IRF3 (green) and +ssvRNA or **(E)** IRF3 (green) and -ssvRNA (magenta), nuclei counterstained with DAPI (blue). Images are 60X confocal microscopy, z-stack max projections, scale bar = 10um, representative of three independent experiments. White asterisk marks cells with nuclear IRF3. **(F)** -ssvRNA puncta were counted per cell, data represent individual cells from three independent experiments with line at mean, **** = p<0.0001 by Mann-Whitney U-test.

To evaluate whether we could detect -ssvRNA during infection of a disease relevant human cell type, we derived mature neurons and glia from human induced pluripotent stem cells (iPSCs). iPSC-derived neural cultures were infected with WNV for 24h and RNAScope for +vRNA revealed high levels of WNV infection across cells (**Fig 6D**). Interestingly, in contrast to A549 cells, far fewer neural cells harbored -ssvRNA, despite replicating +vRNA to high levels (**Fig 6E**). However, the cells that did harbor -ssvRNA accumulated many more puncta than we typically observed in A549s (c.f., **Fig 3J**). Finally, we queried whether accumulation of -ssvRNA in neural cells correlated with innate immune activation and indeed, cells with high levels of -ssvRNA demonstrated nuclear translocation of IRF3 consistent with first responder cells (**Fig 6F**). These data indicate that -ssvRNA accumulation can lead to the activation of IRF3 and innate immune activation in disease relevant human cells and tissues.

## DISCUSSION

Here we define a mechanism of initial innate immune activation during WNV infection to address the paradox of how cytoplasmic sensors detect otherwise sequestered viral RNA. WNV can passively evade innate immune detection at early timepoints by sequestering PAMPs within replication compartments away from cytoplasmic sensors^16^. But as infection progresses, low levels of negative-sense single-stranded viral RNA become exposed in the cytoplasm of infected cells and are recognized by RIG-I to induce innate immune activation. The release of this -vRNA occurs concomitantly with viral particle assembly and is mediated by expression of viral capsid protein within the cell. The accumulation of -ssvRNA varies amongst infected cells, but high levels of -ssvRNA are the defining feature of innate immune activation by first responder cells. Importantly, this pattern of -ssvRNA accumulation and immune activation occurs in disease-relevant cell types both in vitro and in vivo.

Viral replication intermediates have been proposed as RLR PAMPs for many RNA viruses, including WNV^8^. Here we show that indeed RIG-I preferentially interacts with -vRNA over +vRNA in infected cells, despite it being present at 100-1000-fold lower levels than +vRNA. Our findings indicate that RIG-I does not sense WNV +RNA, consistent with the presence of 5’cap structures on the +vRNA that can block RIG-I binding to RNA^11,12^. Therefore, despite high levels of cytoplasmic +vRNA, it cannot act as a PAMP during infection. Our observations instead reveal - ssvRNA as the WNV PAMP that is sensed and bound by RIG-I, causing innate immune activation in the first responder cells. Negative-sense vRNA, is canonically uncapped^43^, making it a much likelier substrate for RIG-I. While we show that -ssvRNA is a substrate for RIG-I, the specific features of this RNA that are bound to RIG-I remain undetermined. In vitro transcription and transfection of segments of WNV vRNA showed that multiple segments of -vRNA were immunostimulatory, including the 3’UTR^44^. The highly structured RNA from this region may serve as a strong PAMP for RIG-I which can sense and bind panhandle structures in RNA^45^. Alternatively, there may exist as-of-yet undefined specific sequences within -vRNA of WNV that stimulate RIG-I, such as the described poly U/UC region in HCV^38^.

Interestingly, transfection of various in vitro transcribed products of the WNV genome revealed that shorter segments of the genome were more potent inducers of innate immunity than full length genomes^44^ reflecting RIG-I’s preference for short RNA substrates^45,46^. Because we used tiled probe sets across the viral genome, we cannot ascertain whether full-length or truncated - ssvRNA is being detected by RIG-I during WNV infection. It is plausible that truncated -ssvRNA products are primary immunostimulatory molecules during infection and that they are generated by cellular enzymes such as XRN1, RNaseL^47,48^ or other molecules. Alternatively, truncated/defective viral genomes that can accumulate during viral RNA replication may act as PAMPs, as has been described for negative-strand virus infection^49^.

Our study shows that exposure of WNV -ssvRNA in the cytoplasm of infected cells is critical for activation of innate immunity. We found that this process occurs at late stages of infection, concomitant with viral egress and particle assembly. During infection, flaviviruses form replication compartments within the ER to sequester replicating RNAs. WNV RCs are exposed to the cytoplasm by a small “neck” that is thought to mediate exchange of nucleotides between the RC and the cytoplasm^14^ and through which nascent RNA must exit to be encapsidated by the capsid (C) protein which mediates assembly into virions at RC proximal assembly sites^50^. The viral NS2A protein is proposed to act as a viroporin to regulate release of vRNA from RCs during flavivirus infection or it may simply facilitate sorting of vRNA into virions^51–53^. Recent studies have shown that during DENV infection, NS2A selectively interacts with 3’UTR of +vRNA to provide selectivity for RNA packaging^53,54^. Here we show that -ssvRNA is released from RCs, raising the possibility that NS2A or other viral proteins might mis-sort or facilitate the release of –ssvRNA into the cytosol where it is then sensed by RIG-I. Indeed, we found that expression of C protein induced the accumulation of -ssvRNA outside of RCs. Capsid proteins bind viral RNA upon egress from RCs and mediate interactions with structural proteins to form virions^55^. The C protein could be involved in regulating release of vRNA from RCs.

The heterogeneous response of cells to viral infection has been observed for infections by multiple viruses across many different families. The heterogeneity in innate immune response can often be linked to the differential kinetics or features of viral replication^22,24,25,56^. Our observations show that there is a wide range in amount of -ssvRNA accumulation amongst individual cells during infection. Thus, the dynamics of –ssvRNA cytosolic accumulation is a driver of differential innate immune activation across a tissue and cell population. Notably, while we find differences in overall levels of -vRNA in innate immune responding cells, there is no difference in +vRNA levels, indicating that differences in genomic and antigenomic vRNA production may occur during viral RNA replication. Differential vRNA production might be controlled by differences in viral genome composition within viral populations or by differences in host factors at an individual cell level. Finally, levels of RIG-I expression within infected cells can influence the capacity of a cell to sense -ssvRNA and induce an innate immune response. In this case, higher levels of RIG-I facilitate sensing of low levels of -ssvRNA within the cytoplasm. Cells that express basally low levels of RIG-I, such as neurons, may require much higher levels of cytosolic accumulation of - ssvRNA to trigger innate immune activation. Even within cell populations, levels of RIG-I are variable^18^, adding an additional factor of complexity to a cells’ ability to initiate an innate immune response.

Overall our study provides new understanding of how RLR-mediated innate immune responses is initiated by flaviviruses. By identifying -ssvRNA outside of replication complexes we provide an explanation for the paradox of how cytoplasmic sensors access viral replication intermediates to trigger innate immune activation. These findings advance our understanding of flavivirus replication and innate immune activation and may provide a generalizable principle that is applicable to other positive strand RNA viruses with similar lifecycles.

## ACKNOWLEDGEMENTS

We thank Nandan Gokhale and Andrew Gustin for critical review of the manuscript. This work is supported by grants from the National Institutes of Health: R01AI145296 (M.G.), F32AI176736, T32AI06677 (E.G.); K08AI150996 (C.S.). The Leica SP8X confocal at the Keck Microscopy Center is supported by NIH grant S10 OD016240.

## AUTHOR CONTRIBUTIONS

Conceptualization, E.G. and M.G.; Methodology, E.G. and C.S.; Investigation, E.G., J.W., J.A., C.S., D.M., and N.E.; Writing – Original Draft, E.G.; Writing - Reviewing and Editing, E.G. J.A., C.S., and M.G.; Supervision, A.O. and M.G.; Funding Acquisition, E.G., and M.G.

## MATERIALS AND METHODS

### Cells and viruses

A549, Vero, B52^57^, and Huh7 cells were cultured in Dulbecco’s Modified Eagles Medium (DMEM) supplemented with 10% fetal bovine serum (FBS), 2mM L-glutamine (Corning), and Penicillin/Streptomycin solution (Corning). HaCaT cells were maintained in DMEM without calcium, supplemented with Peniciillin/Streptomycin solution. A549-IRF3KO cells were described in^58^, A549-RML cells were described in^30^. A549-RIG-I OE (RIG-I overexpression cells) were generated by lentiviral transduction of cells with a selectable pWPI lentiviral vector construct encoding RIG-I (pWPI_RIG-I_blr, a kind gift from Ralf Bartenschlager), selected for transduction with blasticidin 10μg/mL (Invivogen), then single-cell cloned by limiting dilution. All cell lines were routinely confirmed negative for mycoplasma contamination by MycoAlert test kit (Lonza).

West Nile virus strain TX2002-HC (GenBank Accession: DQ176637.1) was plaque purified then amplified in Vero cells. Virus was amplified in Vero cells by infection at MOI of 0.01 for 96 hours in virus growth medium (DMEM, 5% FBS, 2mM L-glutamine, 10mM HEPES (Corning)) and cell supernatant was harvested. Virus titer was determined by plaque assay in Vero cells. The WNV-TX02 infectious clone was used to generate infectious virus as described previously^59^

### Infections

All infections were performed at multiplicity of infection (MOI) of 1.5 plaque forming units(pfu)/cell unless otherwise specified. Virus was diluted in DMEM with 1% FBS and infection was performed at low volume, with rocking at 37°C for 2 hours. Inoculum was then removed and replaced with virus growth medium and incubated for indicated timepoints.

### Mouse infections and tissue preparation

C57BL/6J (B6/J) we obtained commercially (Jackson Laboratories) and bred in house. 8-12 week old mice were used for experiments. Animal studies were carried out under the supervision and approval of the Institutional Animal Care and Use Committee (IACUC) of the University of Washington, protocol number 4298–01 (PI: AO).

For intracranial injections mice were deeply anesthetized. 100 PFU of WNV-TX02 generated from infectious clone were diluted in 20 μl of PBS and injected into the third ventricle of the brain with a guided 29-guage needle. Mock-infected mice were intracranially injected with 20 μl of PBS. At 5 days post infection, mice were anesthetized and perfused with PBS prior to brain dissection. Brain tissue was post-fixed overnight in 4% PFA, followed by cryoprotection in 30% sucrose for 48 h. Brains were then embedded in optimal cutting temperature compound (Tissue-Tek) on the freezing element of a Leica CM3050 S cryostat. 20-μm sections were sliced into 24-well plates containing PBS with 0.05% sodium azide prior to mounting on super frost slides.

### Neural cell cultures

A well characterized human induced pluripotent stem cell line (line CV^60,61^, a kind gift from Jessica Young (University of Washington)) was differentiated into neural progenitor cells (NPCs) using dual-SMAD inhibition techniques as previously described^62,63^. NPCs were maintained in growth medium consisting of DMEM/F12 (Gibco) base, containing: 1x B27 (Gibco), 1x N-2 (Gibco), 1x antibiotic/antimycotic (Gibco), and 1mM L-glutamine (Gibco), supplemented with 20 ng/mL basic fibroblast growth factor (bFGF; Sigma-Aldrich #GF003). Differentiation into mixed neural cells was performed by switching NPCs from growth medium to differentiation medium (DD) for a period of 6 weeks prior to experiments. DD medium consists of DMEM/F12 (Gibco) base, containing: 1x B27 without Vitamin A (Gibco), 1x N-2 (Gibco), 1x antibiotic/antimycotic (Gibco), and 1mM L-glutamine (Gibco), supplemented with 20 ng/mL brain-derived neurotrophic factor (BNDF; Peprotech) and 20 ng/mL leukemia inhibitory factory (LIF; Peprotech). Cells were fed every 2-3 days with fresh medium. After 2 weeks of differentiation, cells were passaged with Accutase (Fisher Scientific) into wells containing glass coverslips #1.5 (Corning) coated with 20 mg/mL poly-L-ornithine (Sigma, P-3655) overnight and 10 mg/mL laminin (Gibco) for four hours prior to plating at 37°C. Cells were differentiated for four weeks on coverslips prior to infection. Infections were performed at low volume with virus diluted in DD media and inoculum was removed and replaced with fresh DD media after 2 hours of infection.

### Flow cytometry and FACS

Cells were fixed in 4% PFA for 15 min at indicated timepoints then permeabilized with methanol on ice for 20 min. Cells were incubated with primary antibody diluted in 1% BSA for 1 hr at RT (anti-Phospho-IRF-3 (Ser386) (E7J8G) XP^®^ Rabbit mAb #37829 (CST) 1:500; anti-dsRNA antibody J2 mouse IgG2a (SCICONS) 1:800; anti-Flavivirus group antigen [D1-4G2-4-15 (4G2)]mouse IgG2a (Absolute Antibodies) 1:1000) washed 2X with PBS, then incubated with secondary antibody diluted in 1% BSA for 30 minutes (anti-rabbit Anti-Mouse IgG2a (γ2a), CF^™^405S antibody produced in goat (Sigma-Aldrich) 1:500; goat anti-rabbit IgG (H+L) highly cross-adsorbed antibody, AlexaFluor 647 1:1000). For flow cytometry cells were analyzed on LSRII (BD). For cell sor)ng, cells were sorted directly into TrizolLS (Invitrogen) on a BD Aria II cell sorter with a minimum of 50,000 cells collected per population. Data was analyzed in FlowJo v9.

### RNA extraction and qPCR

Cells were lysed in Trizol or RLT Buffer and RNA was extracted according to manufacturer’s instructions (Trizol) or with RNeasy Mini Kit (Qiagen) according to manufacturer’s instructions. cDNA was synthesized with iScript Select cDNA Synthesis Kit (Bio-Rad). cDNA was synthesized using a combination of random primer mix and oligo dT provided. qPCR was performed with 1:40 dilution of cDNA, SYBR Green Master Mix (AppliedBiosystems), and forward/reverse primers at 5μM and run on an AppliedBiosystems QuantStudio5. Relative expression of genes was compared to housekeeping gene RPL13a. Primer sequences are listed in **Table 1**.

**Table 1.**
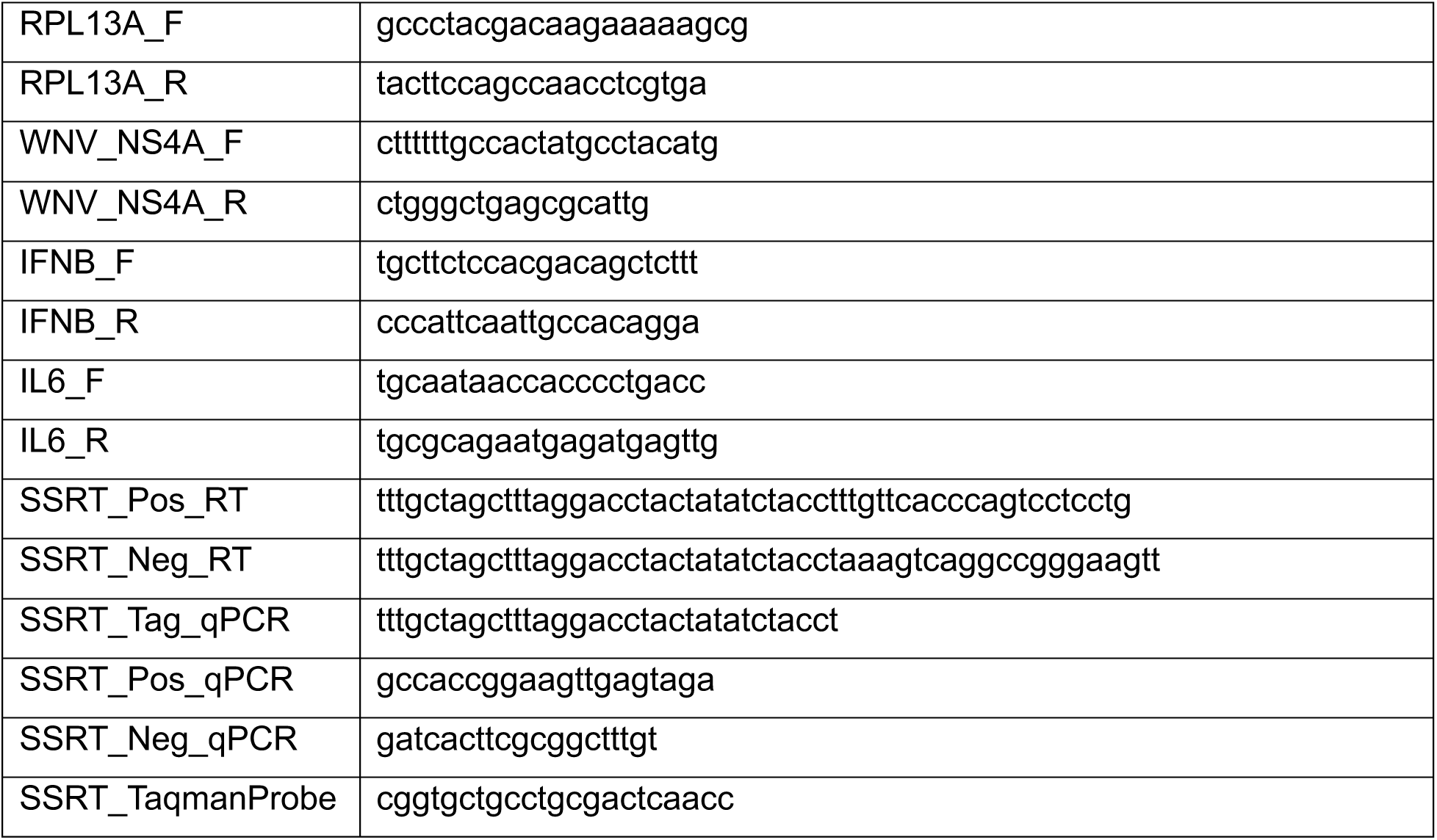
RT-qPCR Primers.

### Strand-specific RT-qPCR

Tag-mediated strand-specific detection of virus was based on a previously described method^31^. In brief, cDNA was synthesized in separate reactions for +vRNA and -vRNA using iScript Select cDNA Synthesis Kit with specific primers and Gene Specific Primer enhancer solution provided, to attach a tag sequence to cDNA. qPCR was performed with 1:10 dilutions of cDNA, 10uM forward/reverse primer, 10uM Probe, and TaqMan master mix (AppliedBiosystems). To generate a standard curve, the region between nucleotides 10,055 and 11,029 was amplified from infectious clone plasmid WNV-CG^59^ and ligated into a pCR-II TOPO vector using a TOPO TA cloning kit (ThermoFisher). The vector was linearized by restriction enzyme digest (HindIII (+vRNA), NotI (-vRNA)) (NEB) and RNA was synthesized by in vitro transcription with T7 (+vRNA) or SP6 (-vRNA) enzymes (Ambion) according to manufacturer’s instructions. Purified RNA was quantified by Qubit Fluorometer (ThermoFisher) and serial dilutions of RNA was reverse transcribed using strand-specific primers for use as standard curve for qPCR. Primer sequences are listed in **Table 1**.

### Transfections

Plasmids were transfected into cells using Lipofectamine 2000 (ThermoFisher Scientific) according to manufacturer’s protocol using a 1:3 ratio of plasmid to lipofectamine. Media was replaced with antibiotic-free media prior to transfection and cells were maintained in antibiotic free media throughout experimentation. WNV Capsid was expressed using pCAGGS with an HA tag on the C-terminus, construction described previously^64^. Control GFP was expressed with pLVX-EIF1α-IRES-Puro containing eGFP (NR-52977, BEI Resources).

HCV polyU/UC RNA was generously provided by HDT-Bio and produced in a GMP facility to ensure quality and purity. RNA (250ng/mL) was transfected using Transit-mRNA transfection kit (Mirus Bio) according to manufacturer’s protocol.

### Replicon generation

A549 IRF3KO cells (4x10^5^ cells) were transfected with 100ng of total RNA extracted from replicon-bearing BHK21 cells^37^ using Transit-mRNA transfection kit (Mirus Bio) with a 1:1 ratio of RNA:Transit-mRNA reagent. Cells were selected for successful transfection at 48h post transfection with media containing 400ug/mL G418 (Fisher Scientific). Selected cells were maintained in 400ug/mL G418 in complete tissue culture media and passaged at no more than 1:3 ratio. Huh7 cells harboring WNV-replicon were described previously^64^.

### Immunofluorescence

Cells were seeded on #1.5 glass coverslips (Corning) prior to infection. At indicated timepoints glass coverslips were washed once in PBS prior to fixation for 15min in 3.7% formaldehyde. Cells were permeabilized with 0.02% Triton-X 100 at RT for 10min. Cells were incubated with primary antibody (anti-NF-κB p65 (L8F6) mouse IgG2b #6956 (CST), 1:500; anti-IRF-3 (D6I4C) XP^®^ Rabbit mAb #11904 (CST) 1:500; anti-dsRNA antibody J2 mouse IgG2a (SCICONS) 1:800) diluted in 1% BSA PBS at RT for 1.5 hrs. Cells were then incubated with secondary antibody (goat anti-rabbit IgG (H+L) highly cross-adsorbed antibody, AlexaFluor 488 1:1000; goat anti-mouse IgG2b cross-adsorbed secondary antibody, AlexaFluor 488 1:1000; goat anti-mouse IgG2a cross-adsorbed secondary antibody, AlexaFluor 594 1:1000) for 1hr at RT then counterstained with DAPI (1:10,000) for 10min a RT and mounted to slides using ProLong Gold antifade mountant.

### RNA-FISH with immunofluorescence

Positive-sense WNV viral RNA was detected using a custom tiled probe set containing 48 probes (**Table 2**) conjugated to Quasar 570 synthesized by LGC Biosearch. Cells were seeded on glass coverslips #1.5 (Corning) prior to infection. At 18 hours post infection cells were washed with PBS then fixed in 3.7% formaldehyde (Fisher Scientific) for 15 min. Cells were permeabilized in 70% ethanol for 1 hr at RT followed by incubation with primary antibody (anti-IRF-3 (D6I4C) XP^®^ Rabbit mAb #11904, CST, 1:500) diluted in 1% BSA in PBS with 160U/mL RNaseOUT (Invitrogen) for 45 min at room temp (RT), then incubation with secondary antibody (goat anti-rabbit IgG (H+L) highly cross-adsorbed antibody, AlexaFluor 488 1:1000) diluted in 1% BSA with RNaseOUT for 40 min at RT. After antibody incubation cells were post-fixed in 3.7% paraformaldehyde for 10 minutes. Cells were washed in wash buffer (2X SSC (Ambion) plus 10% formamide (ThermoFisher) in nuclease-free water) prior to hybridization with FISH probes. Probes were diluted to 5nM in hybridization buffer (wash buffer plus dextran sulfate) and applied to coverslips. Cells were incubated overnight at 37°C in a humidified chamber. Cells were then washed twice with wash buffer for 30 min each at 37°C. On the second wash, DAPI was added to counterstain nuclei. Coverslips were then mounted to slides with ProLong Gold anti-fade mountant (ThermoFisher) and allowed to cure overnight before imaging.

**Table 2.**
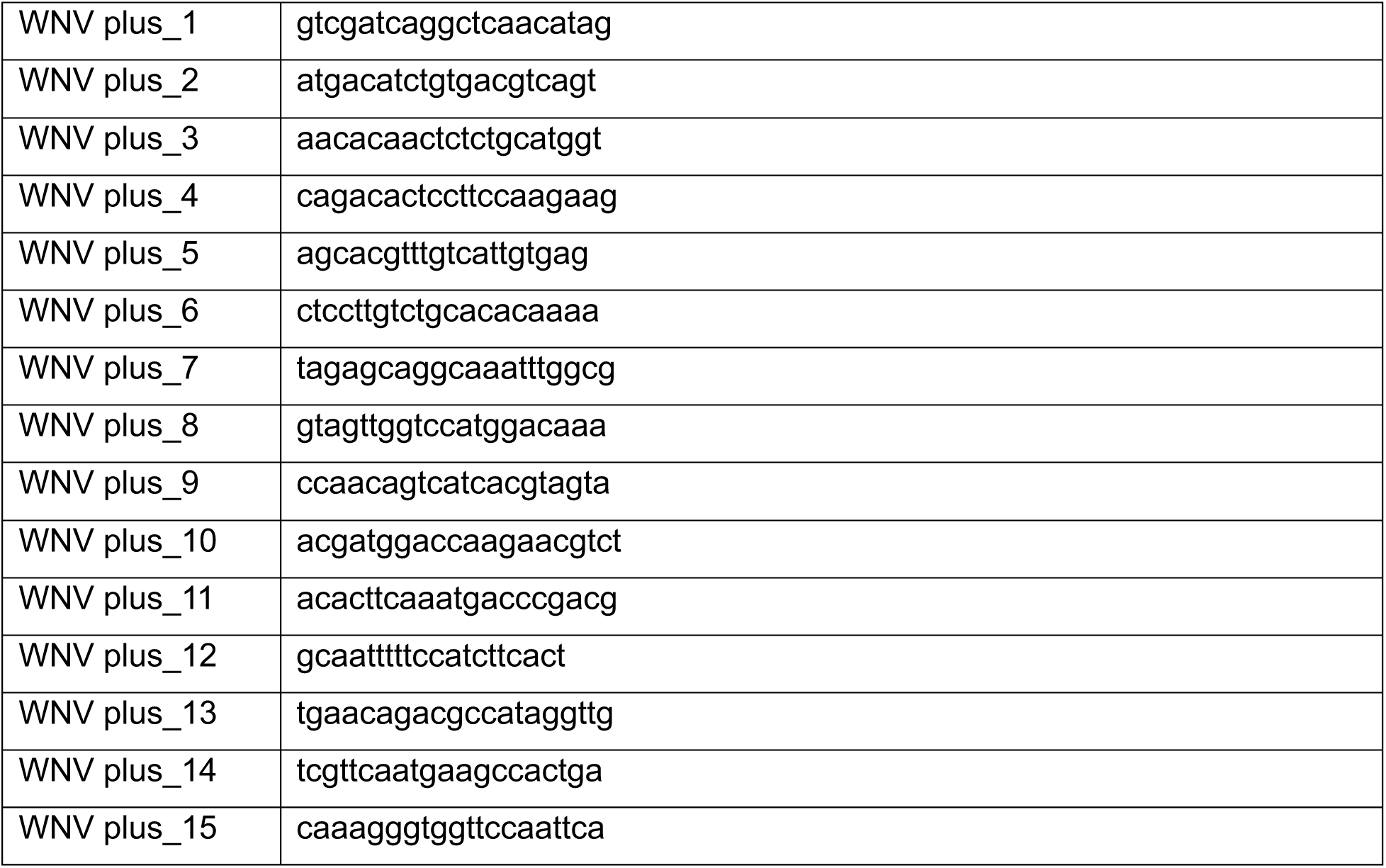
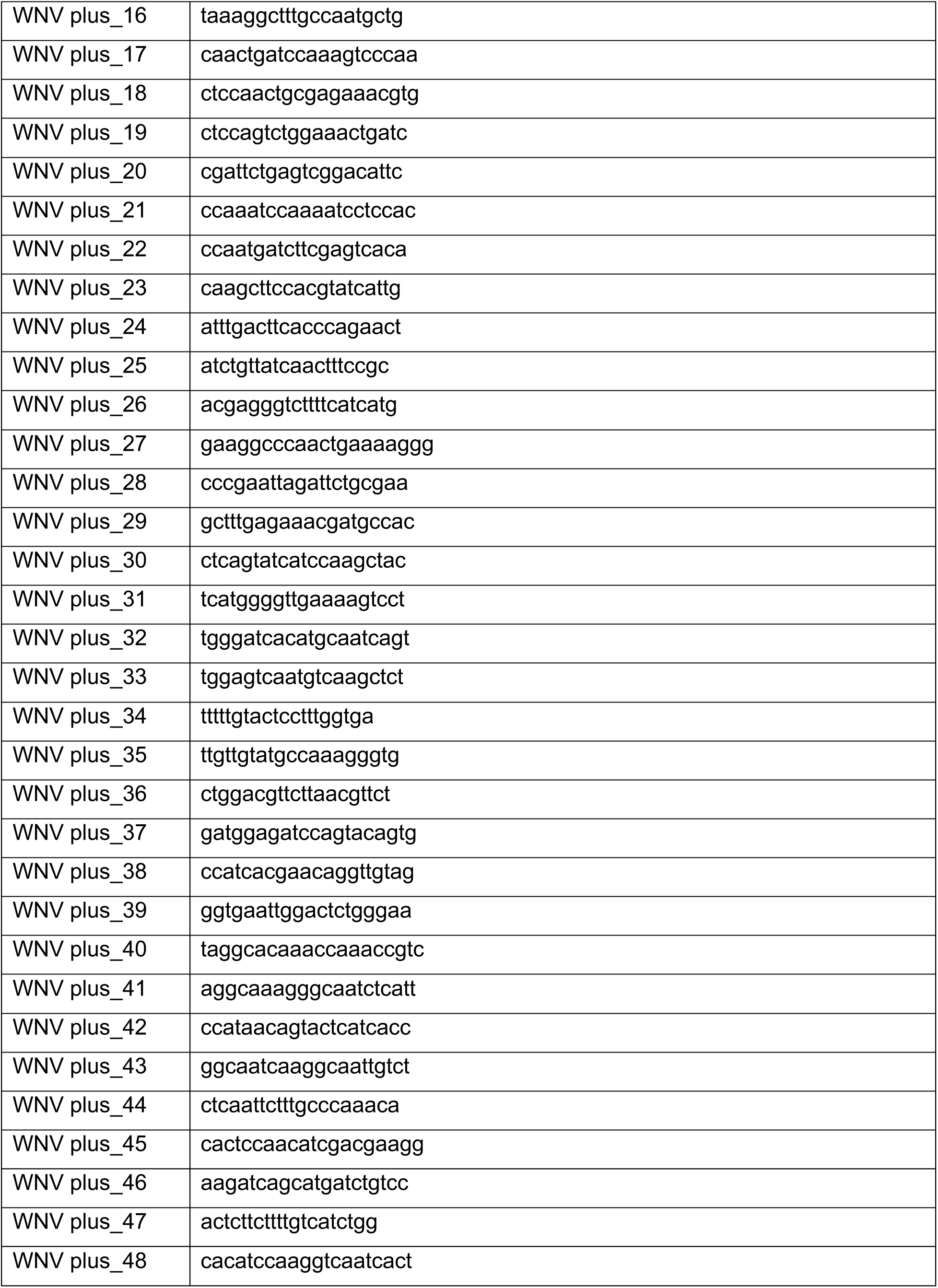
WNV +vRNA RNA-FISH probes.

### RNAscope with immunofluorescence

Positive-sense viral RNA was detected using RNAscope Probe-V-WNV-pp West Nile virus isolate NY99P2 complete genome (Catalog #475091) and negative-sense viral RNA was detected using RNAscope Probe-V-WNV-pp-sense – West Nile virus isolate NY99P2 complete genome (Catalog #553671). Cells were seeded on glass coverslips #1.5 (Corning) prior to infection and fixed at indicated timepoints with 4% paraformaldehyde for 15 minutes at RT. Cells were then washed twice with PBS and subsequently dehydrated into 50%, 70%, and 100% ethanol incubating for 1 min each. Samples were stored in 100% EtOH until further processing. RNAscope was performed with the RNAscope Multiplex Fluorescent Reagent Kit v2 (Catalog #323100, ACDBiotechne) according to manufacturer’s protocol with modifications described below. To begin processing cells were rehydrated into 70% ethanol then 50% ethanol followed by 10 min incubation in PBS. Endogenous peroxidases were quenched by incubating cells in 0.9% H2O2 (w/v) for 10 min at RT then rinsed twice in nuclease-free water before returning to PBS. Cells were then incubated in a humidified chamber with primary antibody (anti-IRF-3 (D6I4C) XP^®^ Rabbit mAb #11904 (CST) 1:500; anti-dsRNA antibody J2 mouse IgG2a (SCICONS) 1:800; anti-Flavivirus group antigen [D1-4G2-4-15 (4G2)]mouse IgG2a (Absolute Antibodies) 1:1000; anti-HA-Tag (C29F4) Rabbit mAb #3724 (CST) 1:500; anti-GFP 9F9.F9 ab1218 mouse monoclonal (abcam), 1:1000; anti-Calnexin (AF18) MA3-027 (Invitrogen), 1:500) diluted in 1% BSA in PBS with 160U/mL RNaseOUT (Invitrogen) overnight at 4°C. The following day cells were washed twice with PBS then post-fixed with 4% paraformaldehyde for 10 minutes at RT. Probes were warmed to 40°C prior to use, then incubated with cells for 2 hours at 40°C and washed thrice with RNAscope Wash Buffer. RNAscope reagents were equilibrated to room temperature then incubated at 40°C in the following sequence with two washes of RNAscope wash buffer in between: Amp1 for 30min; Amp2 for 30min; Amp3 for 15min; HRP-C1 for 15min; diluted Opal 570 for 30min; HRP Blocker for 15min. Opal 570 Reagent (Akoya Biosciences) was diluted 1:3000 in RNAscope TSA Dilution Buffer. Cells were then incubated with secondary antibody (goat anti-rabbit IgG (H+L) highly cross-adsorbed antibody, AlexaFluor 488 (Invitrogen) 1:1000; goat anti-mouse IgG (H+L) highly cross-adsorbed antibody, AlexaFluor 488 (Invitrogen) 1:1000) diluted in 1% BSA with DAPI for 30 min at RT prior to mounting coverslips onto slides with ProLong Gold antifade mountant (ThermoFisher) and cured overnight prior to imaging.

For samples that were denatured prior to probe staining, cells were incubated with 50mM molecular biology grade NaOH (Sigma Aldrich) for 30 sec then rinsed twice with PBS, post-fixation after primary antibody incubation and immediately prior to addition of probes.

For tissues staining, tissue slices were baked on to slides for 60 min a 65°C prior to dehydrating with ethanol. Tissue slides were then processed the same as cells described above with the addition of Protease III Reagent for 40 min at 40°C followed by two rinses with distilled water immediately prior to probe binding.

### Microscopy

Epifluoresence widefield images were acquired on a Nikon Ti Eclipse inverted microscope with a 40X oil-immersion objective (NA 1.3). Confocal images were acquired with a Nikon C2 laser scanning confocal using a 60X oil-immersion objective (NA 1.4). Z-stacks were acquired with 0.25um step size over a 5um Z range and maximum intensity projections are displayed. NS3 and -ssvRNA colocalization images were obtained on a Leica SP8X confocal, 63X oil-immersion objective (NA 1.6). Images were deconvolved using the LASX Expert software lightning deconvolution with global setting.

### Image quantification

Image processing was performed in Fiji (ImageJ2 v2.14.0) with contrast adjustments performed equally across all experiment samples using mock infected cells to set imaging parameters. *Percent Nuclear IRF3* – Cells were binned as nIRF3+ or nIRF3-based on intensity of IRF3 staining that overlaps with DAPI. Infected cells were determined by either staining positive with 4G2 or dsRNA by immunofluorescence or +vRNA by RNA-FISH. Three representative fields of view were quantified per sample with over 150 cells analyzed per experiment.

*Mean intensity of vRNA* – Images were analyzed using a custom built pipeline in Cell Profiler (v4.2.6)^65^. Widefield 40X images were loaded into Cell Profiler. Nuclei were identified by DAPI staining and “Identify Primary Objects” then cytoplasm was identified using Calnexin immunofluorescence staining using “Identify Secondary Objects”. Intensity of vRNA staining was measured on a per cell basis for individually segmented cells. Three representative fields were quantified per sample with over 100 individual cells analyzed per sample per experiment. For measurement of vRNA in nIRF3+ and nIRF3-cells, cytoplasm was identified using “Identify Secondary Objects” based on IRF3 immunofluorescence. Intensity of IRF3 was measured in nuclei and cytoplasm and cells with a ratio > 1 were binned as nIRF3+ and cells with a ratio <1 were binned as nIRF3-.

*Mean dsRNA intensity* - Images were analyzed using a custom built pipeline in Cell Profiler (v4.2.6)^65^. Widefield 40X images were loaded into Cell Profiler. Nuclei were identified by DAPI staining and “Identify Primary Objects” then cytoplasm was identified by expansion based on a standardized micron distance. Intensity of dsRNA staining was measured on a per cell basis for individually segmented cells. Three representative fields were quantified per sample with over 500 individual cells analyzed per sample per experiment.

*-ssvRNA spot counts* – Images were analyzed in Fiji. Maximum projection images were generated from confocal z-stacks. Individual cells were identified with manually drawn regions of interest (ROI’s) based calnexin or IRF3 staining. -ssvRNA channel was thresholded based on mock samples and the “Analyze Particles” function was used to count spots within each ROI. Three to five representative images were analyzed per samples with 20-50 cells analyzed per sample per experiment.

*Mouse tissue* – 60X confocal z-stack images were converted to maximum projection images in Fiji prior to uploading into QuPath for further analysis^66^. Images were segmented using “Cell Detection” based on nucleus parameters with cell expansion to define nuclei. Mean intensity of 4G2 was measured per cell using mock infected animals to define “positive” cells. -ssvRNA were identified using “Subcellular spot detection” using a threshold established based on mock infected tissues. Four representative fields were analyzed per animal with >50 cells analyzed per field.

### RIG-I Immunoprecipitation

Infected A549-IRF3KO cells were trypsinized at 24 hours post infection, resuspended in PBS, and counted. Cells were distributed at a concentration of 1x10^6^ cells/10mL PBS. Formaldehyde was added to each sample to reach a final concentration of 0.3% then incubated with mixing at RT for 10 min. Fixation was quenched by addition of glycine to a final concentration of 250mM, cells were pelleted and washed twice with PBS. Cell pellets were resuspended in 500uL of RIPA buffer [50mM Tris-HCl pH 7.5, 1% NP-40, 0.5% NaDOC, 1mM EDTA, 150mM NaCl] with Halt Phosphatase and Protease inhibitor cocktail (Life Technologies) and 80U/mL RNaseOUT (Invitrogen) and incubated on ice for 20 min. Samples were sonicated with a probe tip sonicator for 30 seconds at 8-9 Watts for 3 rounds with 2 min of resting samples on ice between rounds to prevent overheating. Samples were then cleared by centrifugation at 14000rpm for 20 min at 4°C.

For each sample, 50uL of Protein A Dynabeads (Life Technologies) was prepared by washing twice in RIPA before incubating with 15uL of anti-RIG-I rabbit serum (969) or normal rabbit serum (Life Technologies #31883) diluted in 500uL RIPA Buffer with head-over-tail mixing for 2 hours at 4°C. Anti-RIG-I rabbit serum was generated by immunizing rabbits with a set of pooled 20 amino acid peptides spanning the entire length of the human RIG-I protein. Beads were then washed thrice in RIPA Buffer. Immediately before use beads were incubated with 0.5uL of RNaseOUT per samples for 10 min at RT. Sample was added to beads and incubated with mixing overnight at 4°C. Input (20uL) was saved for each sample. Prior to washing samples, 20uL of unbound lysate was removed to assess efficiency of immunoprecipitation. Samples were washed as follows: 1X with RIPA Buffer; 2X with High Stringency Buffer [20mM Tris pH 7.5, 120mM NaCl, 25mM KCl, 1% Triton-X 100, 0.1% SDS, 1mM EDTA], 3X High Salt Buffer [20mM Tris pH 7.5, 500mM NaCl, 1% Triton-X 100, 1% NaDOC, 5mM EDTA], 3X Low Salt Buffer [20mM Tris pH 7.5, 5mM NaCl, 0.5% Triton-X 100, 5mM EDTA], 2X NT2 Buffer [50mM Tris pH 7.5, 150mM NaCl, 1mM MgCl2, 0.05% NP-40]. After washing, beads were resuspended in 100uL RIPA Buffer. 20uL of sample was reserved for western blot. Samples were reverse crosslinked by heating to 70°C for 45 min while shaking at 1100rpm on a thermomixer to keep beads in solution. Samples were then added to Trizol solution for RNA extraction. Equal volumes of RNA were used for each reverse transcription reaction.

### RNA proximity ligation assay

This assay was performed as described^29^ by modifying the Duolink Proximity Ligation Assay (Duolink In Situ Detection Reagents Orange, Sigma Aldrich). Cells were seeded on coverslips prior to infection then fixed at 18 hours post infection in 4% paraformaldehyde for 15 min, washed twice with PBS, then permeabilized with 0.2% Triton-X 100 for 15 min at RT. Cells were blocked for nonspecific probe binding by incubating in Blocking Buffer [10 mM Tris-acetate, pH 7.5, 10 mM magnesium acetate, 50 mM potassium acetate, 250 mM NaCl, 0.25 µg/µL bovine serum albumin [BSA], and 0.05% Tween 20] with 20ug/mL sheared salmon sperm DNA (Abcam) for 1 hour at 4°C. Cells were then incubated with 100nM probe pools diluted in fresh Blocking Buffer at 4°C overnight. Pools consisted of 6 probes designed to target regions spanning the genome and anti-genome listed in **Table 3**. Cells were then washed thrice in PBS, blocked in PBS with 0.1% Tween-20 with 1% BSA, and 20ug/mL salmon sperm DNA at RT for 1 hr, then washed once with PBS with 2X SSC and 0.1% Tween-20, then once with PBS. Cells were then blocked with DuoLink Blocking Solution for 1 hour at 37°C in a humidified chamber. Primary antibody (anti-RIG-I mouse monoclonal antibody (Alme1), AdipoGen, AG-20B-0009-C100, 1:200) was diluted in Duolink Antibody solution and applied to samples at RT for 1 hr. Cells were washed thrice in PBS then incubated with Duolink In Situ PLA Probe Anti-Mouse MINUS (DUO92004) diluted 1:5 in Duolink Antibody solution for 1 hr at 37°C in a humidified chamber. Ligation, amplification, and labeling were performed according to manufacturer’s instructions. In brief, samples were washed twice with Wash Buffer A for 5 min with shaking. Samples were then incubated with ligation mix for 30 min at 37°C then washed twice with Wash Buffer A for 2 min with shaking. Sample were incubated with amplification mix for 100 min at 37°C then washed twice with Wash Buffer B for 10 min with shaking. On the second wash DAPI was added at 1:10,000 dilution in Wash Buffer B. Cells were then incubated with anti-NS3 antibody (R&D systems MAB2907, 1:500) conjugated to AlexaFluor 647 using the AlexaFluor 647 Antibody Labeling Kit (Life Technologies) diluted in PBS with 1% BSA for 30 min at RT. Coverslips were then mounted on to slides with ProLong Gold anti-fade mountant (ThermoFisher) and allowed to cure overnight before imaging.

**Table 3.**
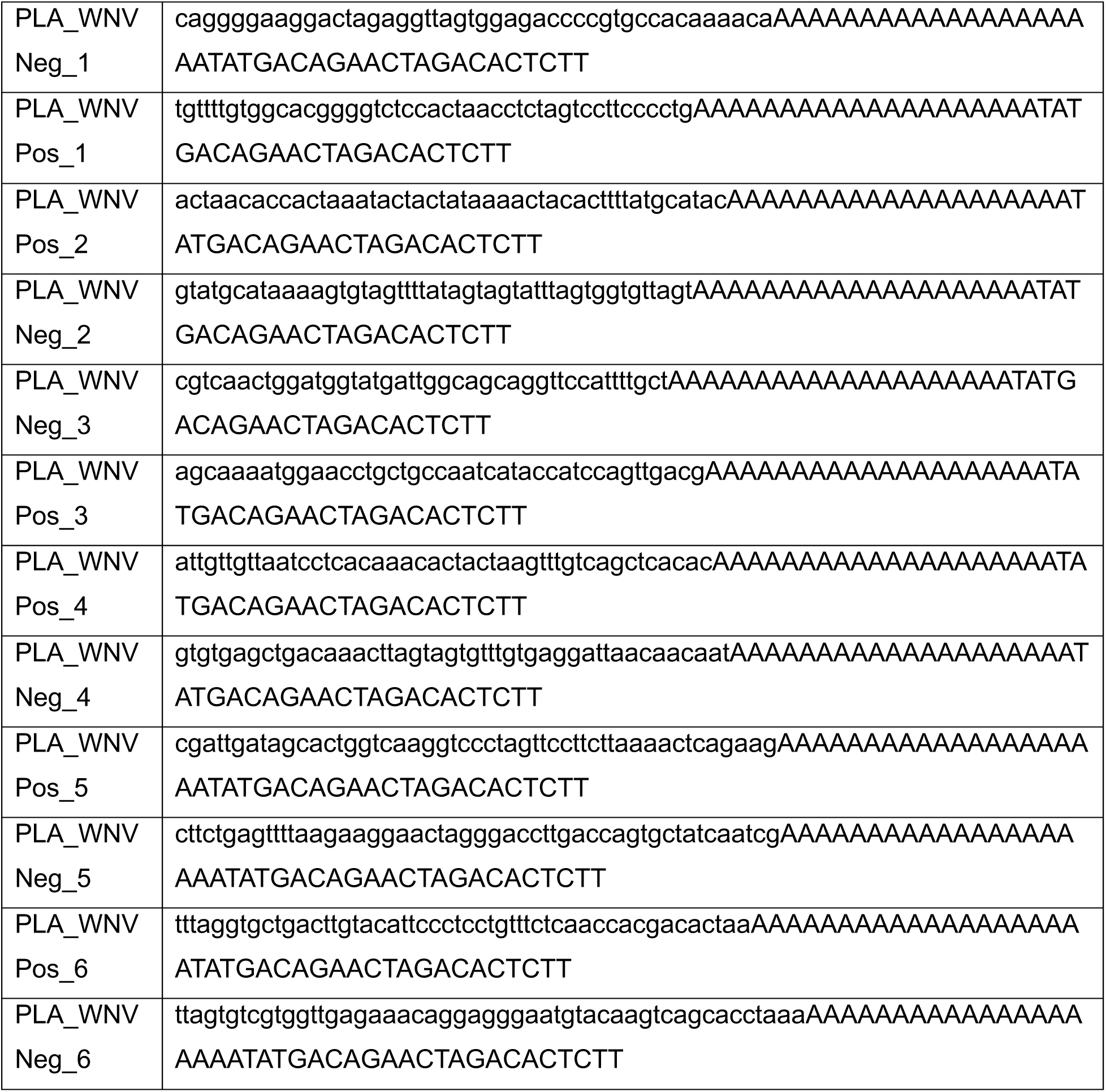
RNA-PLA Probes.

### Western blot

Samples were lysed in RIPA Buffer. Protein concentration was quantified by Pierce BCA Protein Assay Kit (Life Technologies) and equal concentrations of protein was loaded (equal volumes loaded for immunoprecipitated samples). Samples were denatured in Laemli Buffer and boiled for 10 minutes then run on a 4-20% Criterion TGX precast midi protein gel (BioRad) and transferred to a PVDF membrane (Millipore Sigma). Membranes were incubated with primary antibody (anti-RIG-I mouse monoclonal antibody (Alme1), AdipoGen, AG-20B-0009-C100, 1:1000; anti-actin mouse monoclonal antibody clone, Sigma-Aldrich (C4 MAB1501), 1:1000) diluted in 5% BSA in PBST overnight at 4°C. Membranes were then incubated with secondary antibody (goat anti-mouse HRP, Jackson Immunoresearch, 1:10,000) for 1 hour at RT and signal was developed using SuperSignal West Pico Plus chemiluminescent substrate (Life Technologies) and imaged on a BioRad ChemiDoc.

### Permeabilization and RNase Treatment

Cells were seeded on #1.5 glass coverslips prior to infection. At 24 hours post infection coverslips were moved to a fresh plate and washed once with ice cold Treatment Buffer [20mM HEPES pH 7.7, 110mM potassium acetate, 2mM magnesium acetate, 1mM EDTA]. Cells were permeabilized with either 50ug/mL Digitonin (Millipore Sigma), 1% molecular biology grade Triton-X 100 (Fisher Scientific), or DMSO (Millipore Sigma) as vehicle control diluted in Treatment Buffer on ice for five minutes. Permeabilization solution was removed and replaced with appropriate RNase Treatment solution for 30 min at RT. RNase A (ThermoFisher Scientific) was diluted to 100ug/mL in Treatment Buffer. RNase III was diluted to 25U/mL in modified Treatment Buffer [20mM HEPES, pH 7.7, 110mM potassium acetate, 5mM magnesium acetate]. After RNase Treatment cells were fixed with 4% paraformaldehyde for 15 min followed by regular processing for immunofluorescence staining or RNAscope.

### Statistics

Statistics were calculated using GraphPad Prism v10.

## SUPPLEMENTAL FIGURES AND LEGENDS

**Figure S1.**
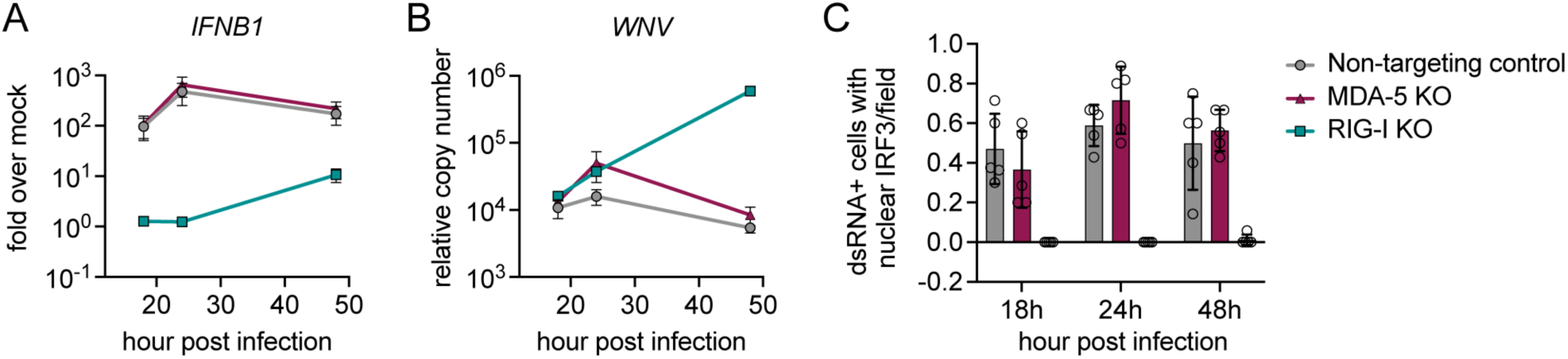
RIG-I exclusively drives an antiviral response during WNV infection in A549 cells. Non-targeting, RIG-I, or MDA-5 CRISPR KO A549 cells were infected with WNV-TX, MOI 1.5, for indicated timepoints and assayed for **(A)** *IFNB1* mRNA and **(B)** WNV RNA accumulation by RT-qPCR. Data is graphed as mean ± SEM for three independent experiments. **(C)** Ratio of nuclear translocation of IRF3 in cells positive for dsRNA was analyzed by immunofluorescence, data represents fields of cells from two independent experiments.

**Figure S2.**
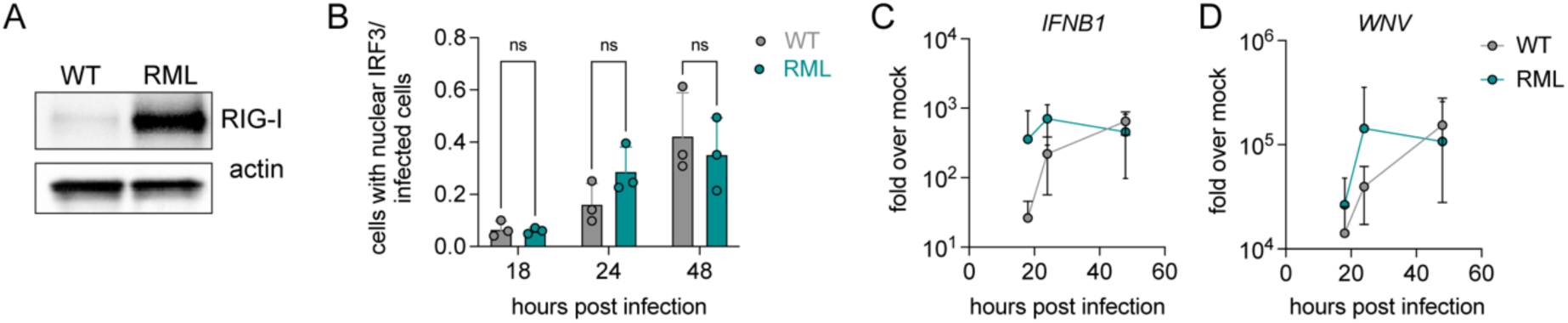
A549-RML have similar innate immune activation kinetics as A549 WT. **(A)** Western blot to detect RIG-I levels in cell lines with actin loading control. **(B)** Infected cells were subjected to immunofluorescence for IRF3 and WNV E protein to detect infection. Ratio of infected cells with nuclear IRF3 is graphed as mean ± SD with individual data points representing average rate of nuclear translocation per experiment, ns = not significant by two-way ANOVA. Indicated cell lines were infected with WNV-TX MOI 1.5 and analyzed by RT-qPCR for **(C)** *IFNB1* mRNA and **(D)** WNV RNA and at indicated timepoints. Data is graphed as mean ± SEM of 3 experiments.

**Figure S3.**
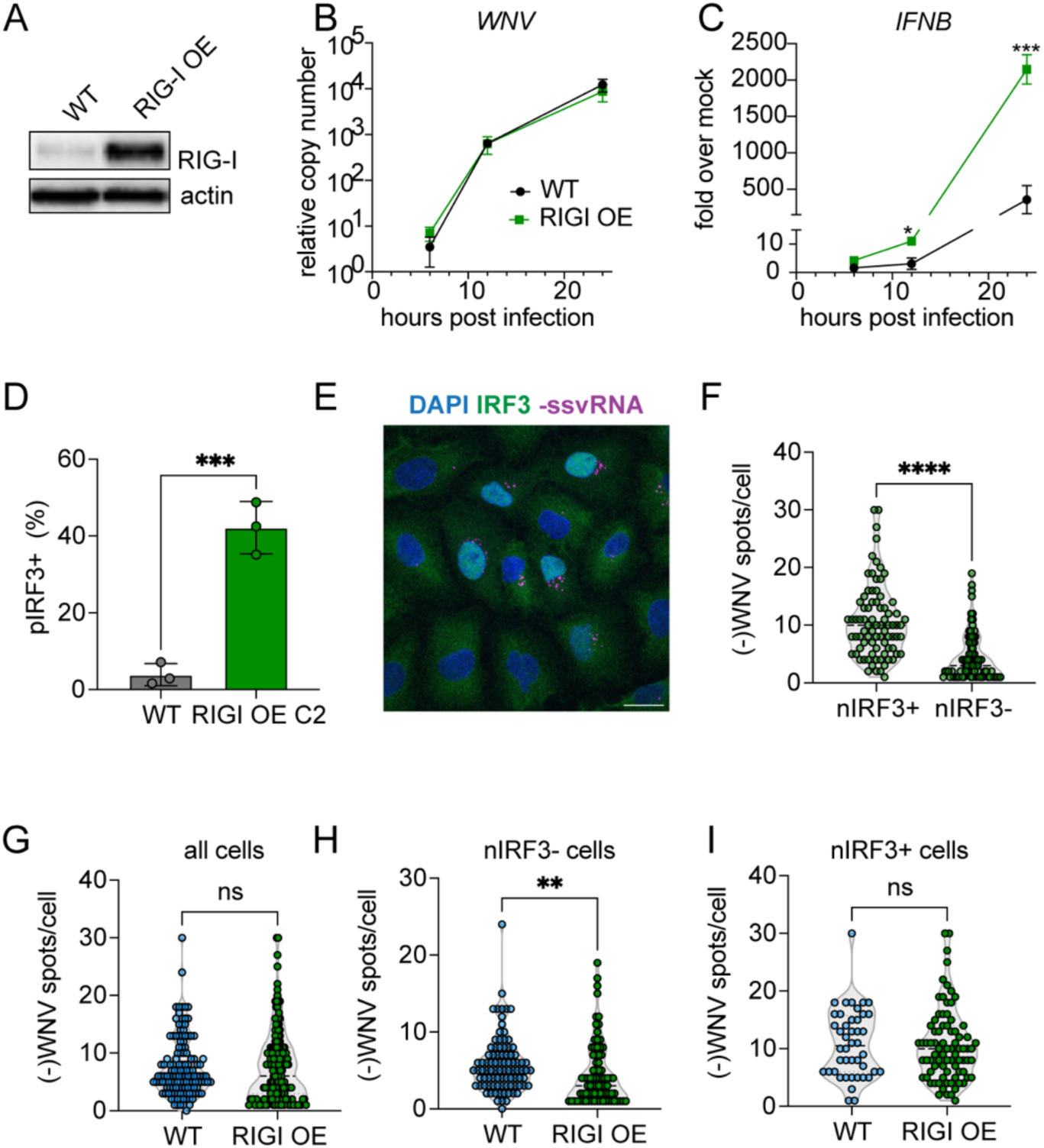
Overexpression of RIG-I lowers -ssvRNA threshold for innate immune activation. **(A)** Western blot to detect RIG-I levels in cell lines with actin loading control. Indicated cell lines were infected with WNV-TX MOI 1.5 and analyzed by qPCR for **(B)** WNV RNA and **(C)** *IFNB1* mRNA at indicated timepoints. Data is graphed as mean ± SEM of 3 experiments, * = p<0.05, *** = p<0.001 by two-way ANOVA with Tukey’s multiple comparisons test. **(D)** Flow cytometry for pIRF3 at 18 hours post infection in indicated cell types, *** = p<0.001 by unpaired t-test. **(E)** A549 RIG-I OE cells were infected with WNV-TX MOI 1.5 for 18h then subjected to combined immunofluorescence for IRF3 (green) and RNAscope to detect -ssvRNA (magenta), nuclei counterstained with DAPI (blue). Image representative of three independent experiments, 60X confocal microscopy, z-stack max projection, scale bar = 20um. **(F)** -ssvRNA puncta were counted per cell and binned based on nuclear IRF3. **(G)** Number of - ssvRNA puncta per cell were compared at 18hpi in all cells in indicated cell types. **(H)** Number of -ssvRNA puncta per cell were compared at 18hpi in cells without nuclear IRF3 **(I)** Number of -ssvRNA puncta per cell were compared at 18hpi in cells with nuclear IRF3. -ssvRNA data displayed as individual cells from three independent experiments overlaid on a violin plot. **** = p<0.0001, ** = p<0.01, ns = non-significant by Mann-Whitney U-test.

